# Substrate reduction therapy in a *Drosophila melanogaster* model of Sanfilippo syndrome

**DOI:** 10.1101/2023.03.27.534462

**Authors:** Sher Li Tan, Laura J. Hewson, Nooramirah Farhana Mustaffar, Qi Qi He, Norbert Wimmer, Paul J. Trim, Barbara King, Marten F. Snel, Kim M. Hemsley, Vito Ferro, Louise V. O’Keefe, Adeline A. Lau

**Affiliations:** Formerly Childhood Dementia Research Group, Hopwood Centre for Neurobiology, Lifelong Health Theme, South Australian Health and Medical Research Institute, Adelaide, SA, Australia, 5000; Currently Department of Molecular and Biomedical Science, School of Biological Sciences, The University of Adelaide, Adelaide, Australia, 5005; School of Chemistry and Molecular Biosciences, The University of Queensland, Brisbane, Australia, 4072; Proteomics, Metabolomics and MS-Imaging Core Facility, South Australian Health and Medical Research Institute, Adelaide, Australia, 5000; The University of Adelaide, Adelaide, Australia, 5005; Currently Childhood Dementia Research Group, College of Medicine and Public Health, Flinders Health and Medical Research Institute (FHMRI), Flinders University, Bedford Park, SA, Australia, 5042

## Abstract

Sanfilippo syndrome, or mucopolysaccharidosis (MPS) types A, B, C or D, are neurodegenerative lysosomal storage disorders resulting from the lack of a specific enzyme involved in heparan sulfate (HS) catabolism. Several treatments are under evaluation for these conditions including substrate reduction therapy, with the most studied compound of this class being the isoflavone genistein. However, recent outcomes from a Phase III clinical trial have shown that high dose oral genistein does not significantly improve neurodevelopmental outcomes in MPS III patients. Here, we have tested an *N*-acetylglucosamine (GlcNAc) analogue inhibitor, 4-deoxy-GlcNAc peracetate, at reducing HS accumulation in cells from patients with Sanfilippo syndrome as a novel substrate reduction therapy. We then confirmed the capacity of this compound to modulate substrate accumulation *in vivo* in a Sanfilippo *Drosophila* model. Treatment with this compound significantly reduced HS in cultured MPS IIIA patient fibroblasts in a time-dependent manner. Neuronal and ubiquitous knockdown *Drosophila* models of MPS IIIC displaying elevated heparan sulfate and behavioural defects exhibited reduced HS burden relative to vehicle-treated controls following oral feeding with the GlcNAc analogue inhibitor. These findings indicate that this compound may be beneficial in slowing the accumulation of HS and may represent a novel therapeutic for Sanfilippo syndrome.

## Introduction

Sanfilippo syndrome, also known as mucopolysaccharidosis (MPS) type III, is a group of inherited lysosomal storage disorders. The classification of sub-type is based on the enzymatic deficiency of a specific lysosomal hydrolase: type A (OMIM#252900; *SGSH*; *N*-sulfoglucosamine sulfohydrolase; EC 3.10.1.1) (1), type B (OMIM#252920; *AAGLU*; *N*-acetyl-α-glucosaminidase; EC 3.2.1.50) (2), type C (OMIM#252930; *HGSAAT;* heparan-α-glycosaminide *N*-acetyltransferase; EC 2.3.1.78 (3), type D (OMIM #252940; *GAS*; *N*-acetylglucosamine-6-sulfatase; EC 3.1.6.14) (4), or type E, though MPS IIIE is yet to appear clinically (5). The estimated birth prevalence varies depending on the sub-type with reports ranging from 1:114,000 to 1:1,407,000 for MPS IIIA and IIIC, respectively, in the Australian population (6).

Given these enzymes are involved in heparan sulfate (HS) glycosaminoglycan degradation, partially-degraded HS fragments accumulate in the tissues and fluids of affected patients eventually causes progressive neurocognitive decline that is clinically similar amongst the sub-types. After an asymptomatic period, Sanfilippo patients exhibit developmental delay, progressing to hyperactivity, aggressive behaviour, loss of social skills and sleep disturbance (7–9). This is followed by severe cognitive deterioration and loss of motor function. Death frequently occurs in the late teens, though the rate of disease progression is variable (10).

Although there are several human clinical trials underway, there is no approved treatment for MPS III. Several mammalian models of MPS IIIA and IIIC exist including murine (11–15) and canine models (16, 17). More recently, *SGSH/CG14291* knockdown *Drosophila melanogaster* models of MPS IIIA have been created to facilitate the study of pathogenic disease mechanisms using genetic screens of candidate genes (18). Characterisation of MPS IIIA *Drosophila* revealed that affected flies share similarities to the human, dog and mouse models including accumulation of HS and motor dysfunction that worsens progressively with age. HS also accumulates in MPS IIIC *Drosophila* following knock-down of the human homologue of human *HGSNAT* (*CG6903*) resulting in behavioural differences in the affected flies (L. Hewson, Honours thesis). Availability of these models permits low cost with high replicate drug, efficacy and toxicity screens to be rapidly carried out in whole organisms.

HS glycosaminoglycans consist of 10-200 repeating disaccharide units composed of D-glucuronic acid (GlcA) 1→4-linked to GlcNAc residues, which are covalently linked to various proteoglycan core proteins via a tetrasaccharide linker (19). The disaccharide repeating unit can be modified to include *N*- and *O*-sulfation and epimerization of GlcA to L-iduronic acid. Previous studies using an analogue of GlcNAc have shown that 4-deoxy-GlcNAc or its peracetylated prodrug form, 4-deoxy-GlcNAc peracetate (4-deoxy-GlcNAc(Ac_3_)), reduce HS levels and chain size in MV3 melanoma or SKOV3 ovarian carcinoma cells (20, 21). HS biosynthesis is also inhibited *in vivo* in zebrafish embryos in a dose-dependent manner (21). Incorporation of 4-deoxy-GlcNAc into the HS chain is not detected and it is hypothesised that the compound may act by decreasing the synthesis of natural UDP-GlcNAc reserves by ‘hijacking’ the cellular machinery to produce UDP-4-deoxy-GlcNAc.

Here, we have evaluated the ability of 4-deoxy-GlcNAc(Ac_3_) at reducing HS accumulation in cells from patients with Sanfilippo syndrome as a potential substrate reduction therapy. We then confirmed the ability of this compound to modulate substrate accumulation *in vivo* in a *Drosophila* models of MPS IIIC.

## Results

### Dose-dependent inhibition of substrate accumulation in cultured MPS IIIA patient cells following 4-deoxy-GlcNAc(Ac_3_) treatment

The experimental compound 4-deoxy-GlcNAc(Ac_3_) (compound **8**) was prepared by a modification of the published methods (**Figure 1**) (22, 23). Full synthetic details and characterization data are provided in the **Supplementary Information**. The parent (or control) compound, GlcNAc peracetate [GlcNAc(Ac_4_)], was prepared by acetylation of GlcNAc (21).

**Figure 1.**
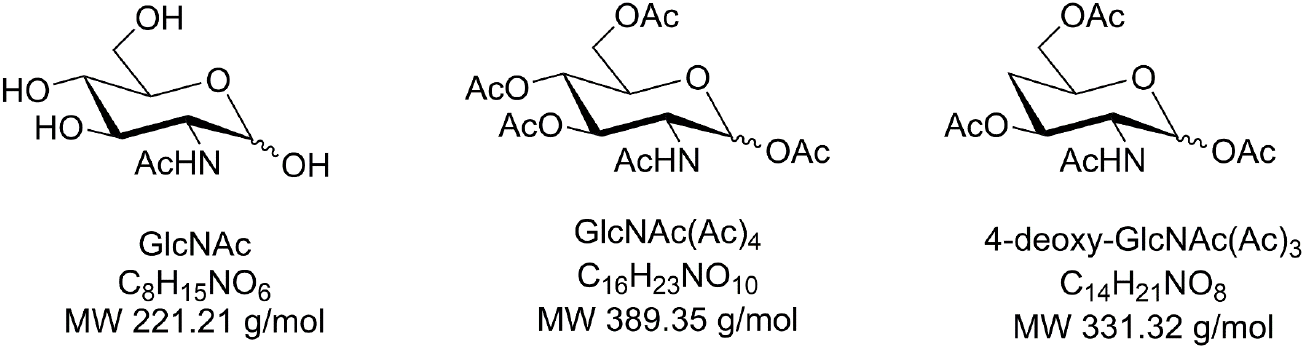
Compound structures. The structures, formulae and molecular weights of *N*-acetyl glucosamine (GlcNAc) and its analogues peracetylated GlcNAc (parent drug/control) and peracetylated 4-deoxy-GlcNAc (experimental drug).

To examine the effect of these compounds on HS biosynthesis, MPS IIIA and apparently healthy control human fibroblasts were treated with escalating concentrations of 4-deoxy-GlcNAc(Ac_3_) or vehicle control for 3 days. The parent GlcNAc(Ac_4_) compound was included as an additional control. Intracellular HS storage in control MPS IIIA cell extracts was 13-fold higher than in water-treated unaffected controls (**Fig. 2a**). HS levels remained consistent in all treated unaffected cells regardless of the compound or dose applied. In contrast, dose dependent effects on intracellular HS were measured in MPS IIIA cells following incubation with 4-deoxy-GlcNAc(Ac_3_), with biologically meaningful reductions in HS only seen at the 16 μM dose. Treatment with GlcNAc(Ac_4_) had no significant effect on HS accumulation.

**Figure 2.**
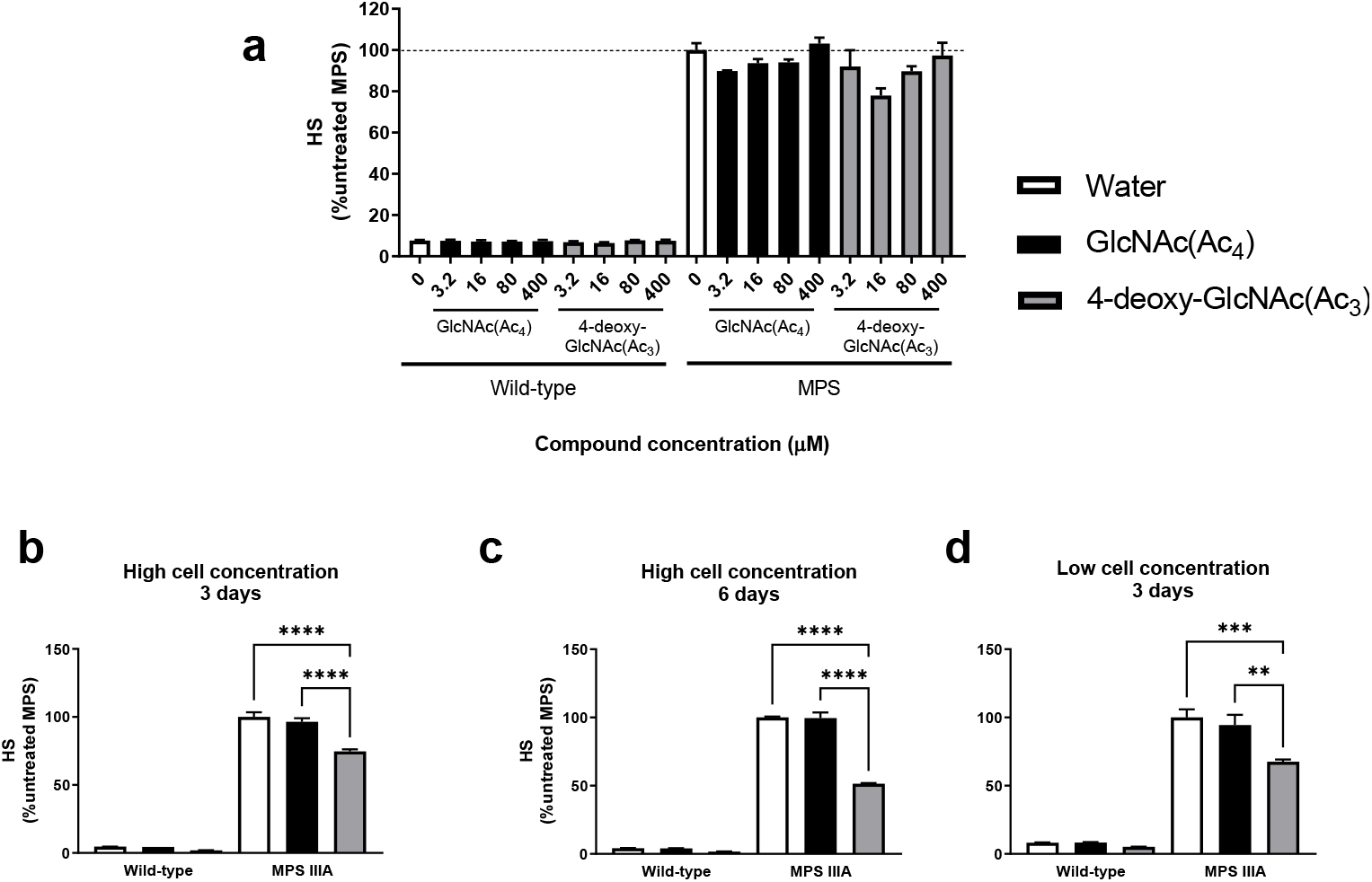
Effect of 4-deoxy-GlcNAc(Ac_3_) treatment on HS in cultured cells. (a) Apparently healthy control or MPS IIIA human skin fibroblasts were plated at 3×10^5^ cells per well and then treated with 4-deoxy-GlcNAc(Ac_3_) at various compound concentrations for 3 days. Water (0 μM) or the parent compound GlcNAc(Ac_4_) at equivalent volumes or compound concentrations were used as controls. HS in cell extracts was measured by tandem mass spectrometry. The level of HS in vehicle-treated MPS IIIA cells is indicated by a dashed line. (b-d) Control or MPS IIIA fibroblasts were plated at 3×10^5^ cells per well (high cell concentration) or 1×10^5^ cells per well (low cell concentration). The effect of either 3 or 6 days treatment with 16 μM of 4-deoxy-GlcNAc(Ac_3_), GlcNAc(Ac_4_) or an equivalent volume of water as a control on HS levels was determined (n=3/group). Data are mean + SEM. ** *p* < 0.01, *** *p* <0.001, \*\*\*\**p* < 0.0001.

Next, we examined the effect of cell density and the duration of treatment. Given that the cells were near confluent (~85-90%) at the initiation of treatment when plated at 3×10^5^ cells per well, we tested whether plating at a lower density of 1×10^5^ cells per well improved the treatment outcomes, as this scenario would see cells actively dividing when the compounds were introduced. As observed in **Fig. 2a**, HS was elevated in the MPS IIIA cells relative to the apparently healthy controls under each treatment condition (**Fig. 2b-d**). Incubation of MPS IIIA cells with 4-deoxy-GlcNAc(Ac_3_) for 3 days reduced HS levels by 25% (high cell dose) or 32% (low cell dose; **Fig 2b,d**). Doubling the length of treatment yielded a 49% reduction in intracellular HS in 4-deoxy-GlcNAc(Ac_3_)-treated cells compared to vehicle-treated controls (**Fig. 2c**). HS levels were similar between MPS IIIA cells incubated with vehicle or parent GlcNAc(Ac_4_) controls.

### HS accumulates in ubiquitous fly models following knockdown of *sgsh* or *hgsnat*

Models of MPS IIIA and MPS IIIC in *Drosophila* have previously been generated using either ubiquitous (actin5c) or neuron-specific (elav2) knockdown of *sgsh/CG14291* or *hgsnat/CG6903* by RNA interference (18; L. Hewson Honours thesis). Comparisons were made against the appropriate wild-type (WT) control line (either actin5c>+ or elav2>+). In actin5c>IIIA flies, *sgsh/CG14291* expression was down-regulated by 71.6% relative to the actin5c>+ control (**Fig. 3a**). *hgsnat/CG6903* expression was likewise reduced in actin5c>IIIC flies (70.6% decrease). HS was quantitated in whole flies at 1 or 15 days post-eclosion by mass spectrometry (24) and found to be significantly elevated in affected flies at both ages, with 3.3-fold and 3.7-fold actin5c>+ HS levels in actin5c>IIIA and actin5c>IIIC flies, respectively at 15 days post-eclosion (**Fig 3b**). No accumulation of HS was detected in whole flies with neuron-specific knockdown in elav2>IIIA and elav2>IIIC lines regardless of age.

**Figure 3.**
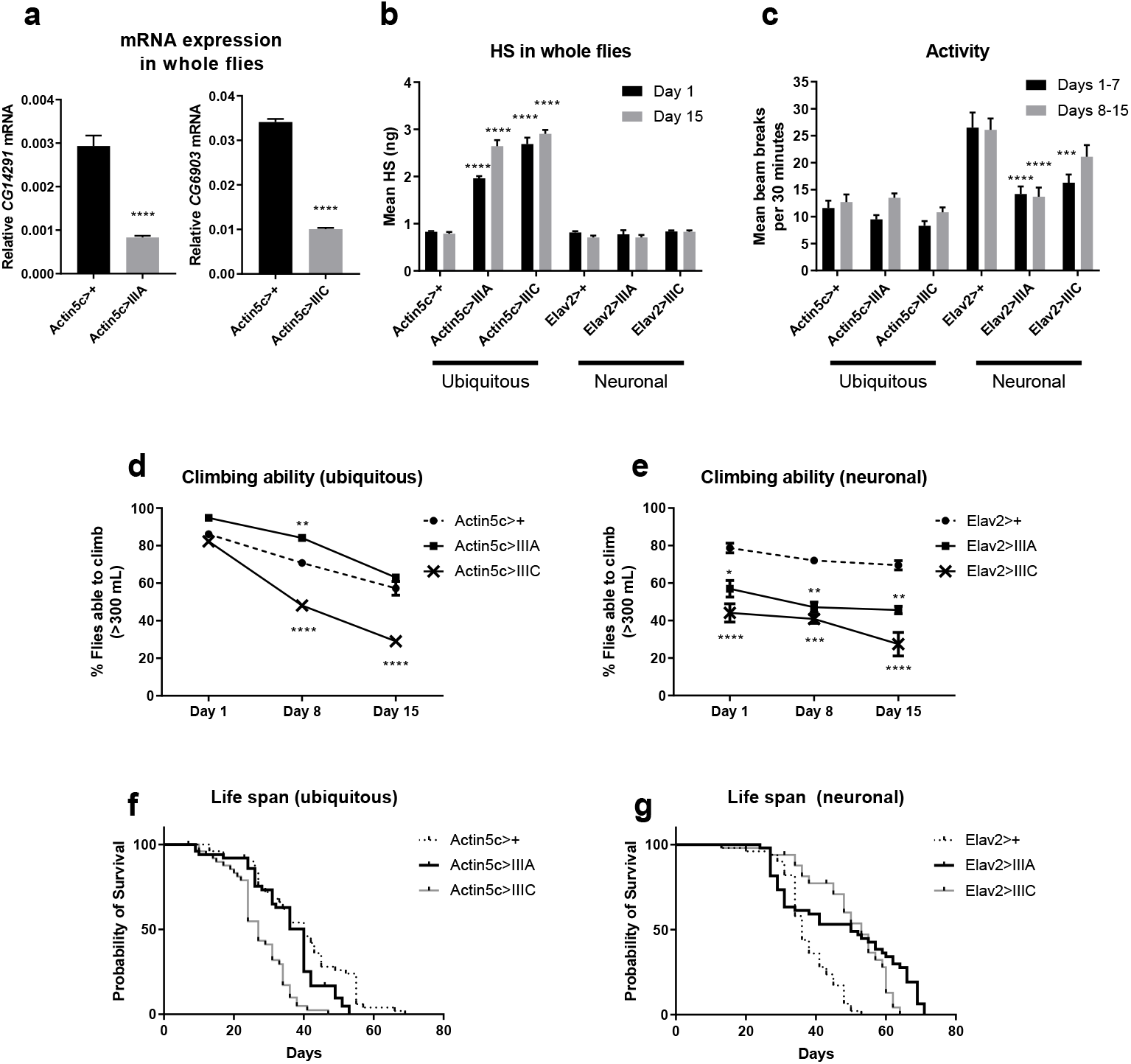
Phenotypic and biochemical characterisation of untreated MPS IIIA and IIIC *Drosophila* lines. (a) Relative quantitation of *sgsh/CG14291* or *hgsnat/CG6903* gene expression compared to the rp49 comparator gene was determined in cDNA prepared from n=6-9 replicates of whole flies. (b) Heparan sulfate (HS) substrate storage in whole flies aged 1-or 15-days post-eclosion was measured by tandem mass spectrometry. n=five biological replicates of five pooled flies per genotype. (c) The average number of beam breaks as an indicator of locomotor activity was determined in male flies individually housed in the *Drosophila* Activity Monitoring system over a 7-day period (n=14-16 males/genotype/time-point; combined lights on and off data). Cohorts of flies were either 1-or 8-days post-eclosion on the first day of recording. (d, e) The negative geotaxis response, classified as an inability to climb above 300 mL in a measuring cylinder, was analysed in the same cohort of male flies at 1-, 8- and 15-days post-eclosion. n=110-179 male flies/genotype/age. (f, g) Kaplan-Meier survival curves were determined in 50 male flies/genotype housed and fed under standard conditions. All data are mean + SEM. * *p* < 0.05, ** *p* < 0.01, *** *p* <0.001, **** *p* < 0.0001 compared to either actin5c>+ or elav2>+ controls.

### Phenotypic and life span deficits in MPS IIIA and MPS IIIC *Drosophila* models

MPS III patients exhibit progressive behavioural abnormalities, including changes in activity, sleep disturbance, the loss of locomotor control and autistic behaviours (7–9). We assessed the disease progression in the untreated affected fly strains to examine the similarity of behaviours to the human clinical phenotype and other MPS III animal models. Beam breaks by flies housed in the *Drosophila* Activity Monitor system were monitored over a 7-day period as a measure of overall activity. Average activities were unchanged in male actin5c>IIIA and actin5c>IIIC flies compared to actin5c>+ at days 1-7 and days 8-15 (**Fig. 3c**; 24 hour data). In contrast, adult elav2>IIIA flies were hypoactive at both time-points. Reduced activity was also exhibited by elav2>IIIC aged 1-7 days, but was not significantly different to controls at ages 8-15 days. The changes in activity were more pronounced in the lights on phase than in the lights off condition (**Supp. Fig. 2a,b**).

The negative geotaxis response was used as a measure of locomotor function in climbing assays. Actin5c>IIIC flies demonstrated a similar climbing ability to actin5c>+ controls at 1-day posteclosion (**Fig. 3d; Supp. Fig. 2c-e**). However, significant and progressive climbing defects were found in flies aged 8- and 15-days, with 49% fewer flies able to climb at the latter time point compared to controls. No significant differences in climbing ability were exhibited in 15-day-old actin5c>IIIA flies compared to controls. Impaired climbing ability was also detected in both elav2>IIIA and elav2>IIIC flies at 1-day post-eclosion and this was maintained at 8- and 15 days post-eclosion (**Fig. 3e**).

MPS IIIA children exhibit decreased social communicative skills akin to autism spectrum disorder behaviours (9) and social interaction in flies can be assessed by measuring the distance from a particular fly to its nearest neighbour (25). No genotype-dependent effects were evident for any of the mutant lines at the time points assessed (**Supp. Fig. 2f,g**).

To determine whether MPS III *Drosophila* models have shortened life expectancy similar to human patients, we conducted lifespan assays (26). In both the ubiquitous and neuron-specific knockdown lines, log-rank (Mantel-Cox) analysis of Kaplan-Meier survival curves showed statistically significant differences between the MPS IIIA or IIIC flies versus the driver-matched control (**Fig. 3f,g**). Whilst the median survival, i.e., the time at which fractional survival equals 50%, was 40 days for both actin5c>+ and actin5c>IIIA, actin5c>IIIC flies had a median survival of 27 days. The maximum survival for actin5c>+, actin5c>IIIA and actin5c>IIIC flies was 60, 46 and 40 days, respectively. In contrast to the actin5c flies, elav2>IIIA and elav2>IIIC flies had unexpectedly significantly longer median survival times than elav2>+ flies: the median survival for elav2>IIIA and elav2>IIIC was 50 days and 53 days, respectively, compared to 36 days for elav2>+ controls. The maximum survival of elav2>+, elav2>IIIA and elav2>IIIC flies was 50, 70 and 63 days, respectively.

### Determination of compound concentration for oral delivery to *Drosophila*

Considering that 4-deoxy-GlcNAc(Ac_3_) slowed the accumulation of intracellular HS in Sanfilippo cell models, we next aimed to determine whether this compound could likewise inhibit HS biosynthesis *in vivo* following treatment in Sanfilippo *Drosophila*. Presuming equivalent metabolism and toxicity of GlcNAc, a dose-response curve was undertaken to identify the highest concentration of GlcNAc that was non-toxic to *Drosophila*. Supplementation of the highest dose of 100 mM GlcNAc to food led to a significant shortening of lifespan in actin5c>+ flies (9 days compared to control flies fed normal food; **Supp. Fig 3**). Statistically significant differences in survival were not evident at 67 mM GlcNAc, the next highest dose assessed, compared to the vehicle-supplemented controls and thus this concentration was used in the subsequent studies.

### Oral delivery of peracetylated 4-deoxy-GlcNAc(Ac_3_) inhibits HS accumulation in MPS IIIC *Drosophila*

We elected to use the ubiquitous actin5c>IIIC fly model to evaluate the ability of 4-deoxy-GlcNAc(Ac_3_) in preventing the intracellular storage of HS as they expressed both behavioural deficits (climbing assay and hypoactivity) and stored quantifiable levels of HS. Actin5c>IIIC flies were treated from hatching with 67 mM 4-deoxy-GlcNAc(Ac_3_), GlcNAc(Ac_4_) or GlcNAc or an equivalent volume of water diluted in normal food media as a control. Significant genotype effects were found with elevated HS measured in all actin5c>IIIC treatment groups compared actin5c>+ controls regardless of age or drug treatment (**Fig 4a**). At 1-day-post-eclosion, actin5c>IIIC flies receiving the control diet (i.e. water-supplemented group) already had a HS burden that was ~3.3-fold more than age-matched actin5c>+ control levels. Significant age effects on HS were also evident, with progressive increases in HS storage measured. *Post-hoc* testing revealed that there was significantly less HS in actin5c>IIIC flies fed 4-deoxy-GlcNAc(Ac_3_) for 15 days as compared to treatment with the GlcNAc(Ac_4_) parent compound over the same time frame (27% less).

**Figure 4.**
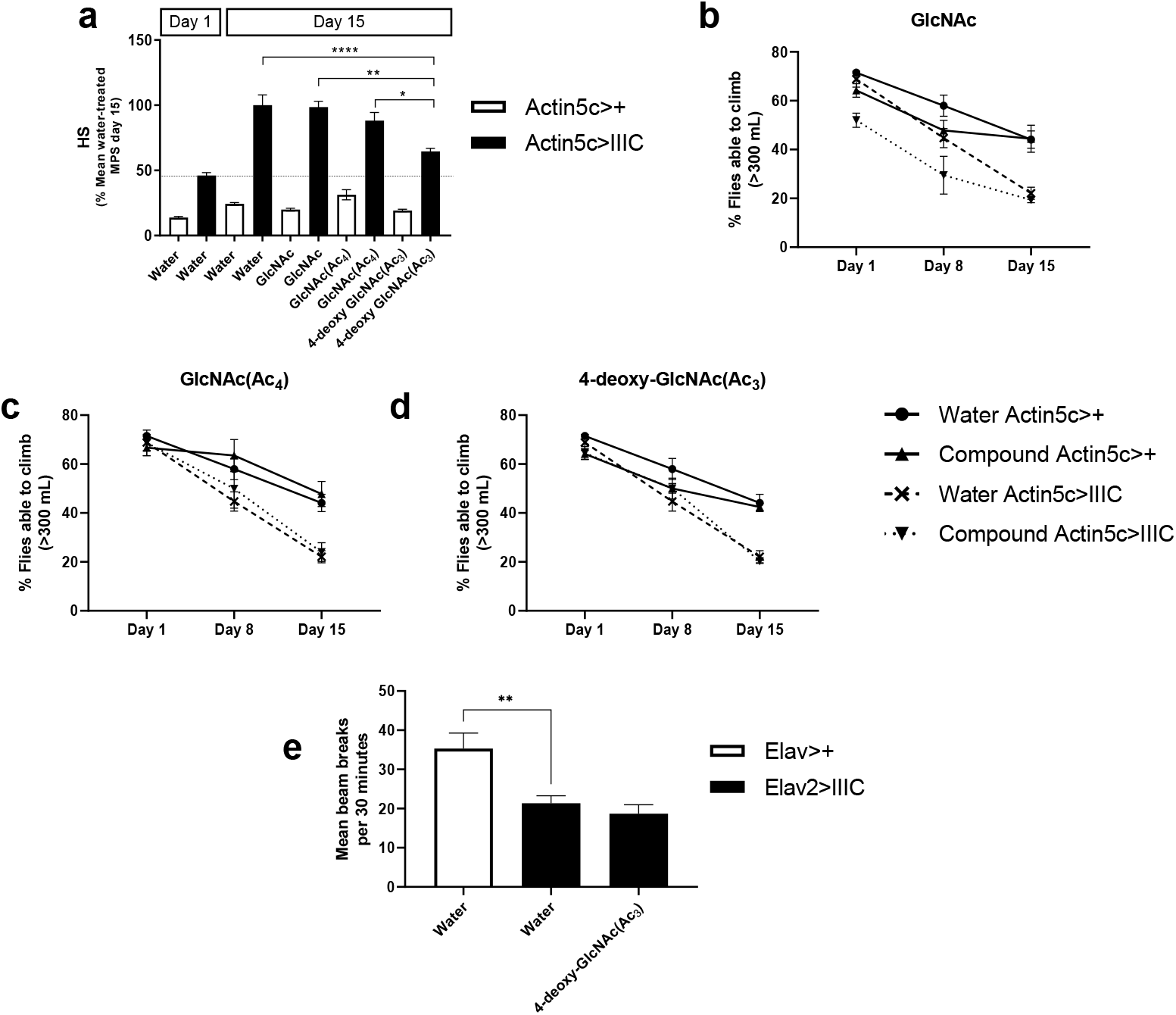
Effect of substrate reduction therapy on HS substrate levels and behaviour in MPS IIIC flies. (a) HS was measured after feeding of 67 mM 4-deoxy-GlcNAc(Ac_3_) in food agar from eclosion for 1 or 15 days. Control diets included an equivalent volume of water, or 67 mM GlcNAc or 67 mM GlcNAc(Ac_4_). Samples were n=5-10 biological replicates of five actin5c>+ (open bars) or actin5c>IIIC (filled bars) flies per group. The dashed line represents the level of substrate storage in untreated actin5c>IIIC flies at the initiation of treatment. (b-d) The effect of supplementing food agar with (b) 67 mM GlcNAc, (c) 67 mM GlcNAc(Ac_4_) or (d) 67 mM 4-deoxy-GlcNAc(Ac_3_) on climbing ability was measured using climbing assays on the same cohorts of male flies at 1-, 8-or 15-days posttreatment. Flies were defined as able to climb if they climbed higher that the 300-mL mark of a measuring cylinder. n=94-179 flies. (e) Elav2>IIIC flies were fed with 67 mM GlcNAc(Ac_4_) in food agar for 10 days post-eclosion and before activity was measured in a Drosophila Activity Monitoring system for 7 days. n=13-16 male flies per group. Data are mean + SEM. * *p* < 0.05, \*\**p* < 0.01, **** *p* < 0.0001 compared to either actin5c>+ or elav2>+ controls.

Given that the HS burden was improved in actin5c>IIIC flies orally supplemented with 4-deoxy-GlcNAc(Ac_3_), we further tested the ability of this compound to prevent the development of behavioural symptoms in affected flies. All groups of flies had comparable climbing ability regardless of genotype or dietary modification at 1-day post-treatment (**Fig 4b-d**). The overall climbing performance was poorer in the affected actin5c>IIIC flies compared to actin5c>+ flies and in older flies compared to either of the younger ages (genotype and age effects of *p*<0.000). However, whilst there were significant differences between the groups depending on the diet on which the flies were raised (treatment effect of *p*<0.002), there was no difference in the motor phenotype between the flies treated with 4-deoxy-GlcNAc(Ac_3_) and any of the three control diets (all *p*>0.05). Likewise, elav2>IIIC flies raised on a diet of 4-deoxy-GlcNAc(Ac_3_) for 10 days post-eclosion remained hypoactive similar to control affected flies when their behaviour was quantitated in an activity monitor (**Fig 4e)**.

To examine whether earlier initiation of therapy could prevent behavioural defects from appearing, flies were crossed directly onto food agar supplemented with 67 mM 4-deoxy-GlcNAc(Ac_3_). However, only one male actin5c>IIIC fly hatched under these conditions (3.45% survival; data not shown) and so further testing could not occur.

## Discussion

Defects in the degradation of glycosaminoglycans results in 11 distinct MPS disorders, each due to the lack of a specific lysosomal enzyme. For some of these conditions such as MPS IVA and VI, and the non-neuropathic forms of MPS I and II, treatments exist in the form of enzyme replacement therapy where the recombinant enzyme is infused intravenously on a regular basis, i.e., Vimizim™, Naglazyme^®^, Aldurazyme^®^ and Elaprase^®^ (27). Intravenous injection of the recombinant enzymes at conventional doses is insufficient to effectively treat neuropathology due to the inability of the enzyme to efficiently traverse the blood-brain barrier. Intra-cerebrospinal fluid infusion or modification of the enzyme with tags to facilitate delivery into the central nervous system is possible, but these modalities will require repeated delivery of enzyme, usually in a clinical setting. Gene therapy vectors expressing lysosomal genes for the treatment of MPS sub-types are also under evaluation in clinical trials with these studies delivering the vector particles directly into the brain parenchyma or intravenously (e.g. clinical trials NCT03612869, NCT02716246). Pre-existing immunity against viral capsid proteins will exclude some patients from viral gene therapies (28). Both central nervous system-directed enzyme replacement and gene therapy carry the risk of potential antibody-related responses to the enzyme and/or vector.

To address these issues, we have investigated substrate reduction therapy with 4-deoxy-GlcNAc(Ac_3_) which aims to reduce HS synthesis and thus slow down the build-up of substrate using affordable, small molecule drugs that can cross the blood-brain barrier. SRT may re-dress the balance between the substrate burden and enzyme activity for those patients with residual active enzyme or slow substrate accumulation in those without active enzyme. The viability of this approach is demonstrated by the availability of FDA-approved glycosphingolipid substrate inhibiting drugs for the lysosomal storage disorders, Gaucher Disease type I and Niemann-Pick type C (e.g., Cerdelga^®^, Zavesca^®^).

At least three compounds have been evaluated for MPS III as potential drugs that interfere with HS biosynthesis. Suberanilohydroxamic acid (SAHA or Vorinostat) is a pan-histone deacetylase inhibitor that was identified by screening of a chemical library for inhibition of the HS modifying enzyme *N*-deacetylase/*N*-sulfotransferase (NDST1) (29). Treatment of MPS IIIA and IIIC patient fibroblasts with SAHA inhibited glycosaminoglycan biosynthesis (29). Compounds rhodamine B and genistein also slow HS/glycosaminoglycan accumulation and improve MPS IIIA or IIIB mouse behaviour (30–32). Translation of genistein treatment into patients produced variable neurocognitive outcomes. A trial of 5 mg/kg/day genistein appeared to improve cognition in an open-label study (33). However, an open-label trial using a much higher dose (150 mg/kg/day genistein) was unable to demonstrate a consistent decline in urinary glycosaminoglycan levels, and after one year of treatment MPS III patient symptoms were either unchanged (89%) or further declined (11%) (34). These results were confirmed in a double blinded, randomised and placebo-controlled Phase III trial in MPS IIIA patients who were given 160 mg genistein/kg/day (or placebo) for one year, followed by an additional year of open label genistein at the same dose (35). Whilst genistein was found to be safe, no apparent clinical benefit was conferred.

Here, we have demonstrated a slowing of intracellular HS accumulation with 4-deoxy-GlcNAc(Ac_3_) delivery as a potential substrate reduction therapeutic approach in cells from patients with MPS IIIA. Further, MPS IIIC *Drosophila* who were fed any of the three control diets progressively accumulated HS. In contrast, significantly less HS was found in MPS IIIC flies fed 4-deoxy-GlcNAc(Ac_3_)-supplemented food for 2 weeks from eclosion compared to age-matched MPS IIIC flies. Despite these decreases in HS biosynthesis, climbing ability and hypoactivity were not improved in 4-deoxy-GlcNAc(Ac_3_)-treated MPS IIIC flies. One explanation for the lack of behavioural improvement may be because the drug treatment was initiated at hatching (eclosion), an age where untreated MPS IIIC flies already exhibited ~3.3-fold more HS than wild-type flies. Previous studies in an MPS IIIA mouse model showed that intra-cisternal enzyme replacement therapy reduces HS more effectively when initiated in young mice compared to older, symptomatic mice (36). This is further supported by the outcomes of a Phase I/II clinical trial in MPS IIIB patients receiving intracranial injections of recombinant adeno-associated viral vector serotype 2/5 vector encoding *NAGLU*. Of the four patients treated in this trial, the youngest child (who was 20 months at the inclusion in the trial) showed the greatest neurocognitive development over the duration of the study whereas the oldest patient at inclusion (53 months) showed the least improvement in developmental quotient (37).

To evaluate whether earlier initiation of treatment could positively impact behaviour, we provided MPS IIIC flies with 4-deoxy-GlcNAc(Ac_3_) throughout the larval stages. However, the compound had a significant impact on development, with flies failing to efficiently hatch. HS plays a critical role in development, cell homeostasis and disease (reviewed in 38). van Wijk and colleagues (21) demonstrated that cell surface HS chain size is reduced following 4-deoxy-GlcNAc(Ac_3_) treatment, which impacted angiogenesis due to the altered binding capacity of HS to fibroblast growth factor-2 and vascular endothelial growth factor. Further, genetic knockdown of HS biosynthetic enzymes sulfateless (sfl) or slalom (sll) in MPS IIIA *Drosophila* worsened the climbing defect that was present in the affected flies (18). Future studies titrating the administered drug dose may be beneficial, where the compound is supplemented early in the disease course from mating, but at a reduced dose throughout larval development.

A secondary outcome of this study was the side-by-side characterisation of two *Drosophila* Sanfilippo models. Elevated HS was only detected in the ubiquitous actin5c-knockdown Sanfilippo models. Despite our inability to detect elevated HS levels in the neuronal knockdown Elav2>IIIA and Elav2>IIIC lines, behavioural differences were apparent compared to the Elav2>+ controls and the outcome in climbing assays reflects that seen in Webber et al. (18) where impaired negative geotaxis responses were found. The negative geotaxis response progressively worsens with age in MPS IIIA mice (39), along with the development of other motor deficits such as neuromuscular grip strength (14, 39). This is consistent with descriptions of motor function deterioration in MPS IIIA and IIIC children where patients eventually lose the ability to walk without support (7, 8, 40, 41). Surprisingly, climbing defects were not found in actin5c>IIIA flies and further experiments are required to elucidate why this innate escape response is unchanged in this line. Another clinically-relevant behavioural change that was identified in this study was the reduced activity of Elav2>IIIA and Elav2>IIIC flies in the *Drosophila* activity monitoring system. Open field hypoactivity has also been described in MPS IIIA and MPS IIIC mice (42, 43) and this may correspond to the later, quieter phase in Sanfilippo children as they become bed-ridden. Changes in the frequency of beam breaks as a measure of activity were not evident in actin5c>IIIA or actin5c>IIIC models and may relate to the low level of overall activity in the actin5c>+ controls.

Significant differences in Kaplan-Meier survival curves were found in ubiquitous knockdown actin5c>IIIA and actin5c>IIIC *Drosophila* compared to the actin5c>+ controls. This correlates with reports of Sanfilippo patients having reduced life spans, with mean survival ages between 13 and 18 years (MPS IIIA) and 19 to 34 years (MPS IIIC) (reviewed in 44). In contrast, both neuronal knockdown models of MPS IIIA and IIIC lived longer than Elav2>+ controls. We attribute this to the potential toxicity of the unbound Gal4 transcription factor in the wild-type line, as has previously been described in neurons and in the developing eye (45, 46). In the GAL4/UAS system employed here, the Gal4 protein binds to the upstream activation sequence to drive the expression of the RNAi to knockdown expression of the lysosomal gene homologue of interest. However, in the control lines, the Gal4 is unbound and can lead to loss of neurons, potentially through apoptotic pathways (46).

In summary, we have demonstrated that oral dosing of 4-deoxy-GlcNAc(Ac_3_) significantly slows the accumulation of HS in a *Drosophila* model of MPS IIIC. Further tests in other model systems are needed to determine whether this level of reduction in HS (27%) is sufficient to mediate phenotypic improvements on its own in the absence of adverse effects. This substrate reduction therapy approach may provide a synergistic benefit when applied as a combination therapy with other treatment approaches. Proof-of-concept studies by Lamanna *et al*. (47) have established that less exogenous recombinant SGSH enzyme is required to correct HS levels in MPS IIIA mice when combined with genetic reduction of HS biosynthesis than when applied as an enzyme replacement strategy alone. Should further testing of this small molecule drug in mammalian models of MPS III continue to reveal that it is efficacious and safe, this compound has potential as a substrate reduction therapy for not only MPS III, but for any MPS disorder that stores HS (MPS I, II, IIIA-D, and VII).

## Materials and Methods

### Compound synthesis

GlcNAc, (*N*-acetyl-D-glucosamine) was obtained from Carbosynth Limited, Berkshire, UK. Peracetylated GlcNAc, referred to as GlcNAc(Ac_4_), was synthesised by acetylation of GlcNAc with excess acetic anhydride in pyridine (21). The synthesis of peracetylated 4-deoxy-GlcNAc (**8;** 4-deoxy-GlcNAc(Ac_3_), C_14_H_21_NO_8_; 331.318 g/mol) was performed as shown in Scheme 1 (**Supplementary Fig. 1**) by a modification of previously described synthetic routes (22, 23). Full details are provided in the **Supplementary Information**. In brief, GlcNAc (**1**) was converted into its methyl-α-glycoside (**2**) via Fischer glycosidation. Selective benzoylation gave the alcohol **3** which was then deoxygenated via conversion to the chloride **4** and treatment with tributyltin hydride to give the 4-deoxy derivative **5**. Saponification of the benzoate groups and acetylation gave **7**. Finally, acetolysis of the methyl glycoside with acetic anhydride and concentrated sulfuric acid gave the target compound **8** as a mixture of anomers.

### Cell culture

Human fibroblast cell lines were obtained from the NIGMS Human Genetic Cell Repository at the Coriell Institute for Medical Research (GM00038 apparently healthy control and GM00629 MPS IIIA). Cells were maintained in Minimum Essential Medium Eagle with Earle’s salts (Sigma M5650) containing 15% (v/v) FBS (Life Technologies 10099-141), 1% (v/v) GlutaMAX (Life Technologies 35050-061) and 1x Penicillin-Streptomycin (Life Technologies 15070063) at 37°C with 5% carbon dioxide. Cells were seeded at a density of 1×10^5^ or 3×10^5^ in 6-well trays and allowed to attach overnight. Compounds at final concentrations ranging from 3.2 to 400 μM in media (or an equivalent amount of water as a vehicle control) were then applied to the cells and cultured for 3 days, or 6 days with media containing the drug replenished on day 3. The cells were detached with 1x trypsin-EDTA (Life Technologies 15400-054), centrifuged and the cell pellet washed once in PBS. After centrifugation, the cells were resuspended in 200 μL of 20 mM Tris, 500 mM NaCl, pH 7.2 and the intracellular contents were liberated by sonication (20 seconds, 30% power; model CV188 Vibra-Cell; Sonics and Materials). The total protein concentration was determined using a Micro BCA™ Protein Assay Kit (ThermoFisher Scientific 23235).

### *Drosophila* lines and crosses

P{GAL4-elav.L}s2/CyO (8765, referred to as elav2,) and P{Act5c-GAL4}25FO1/CyO (4414, referred to as actin5c) were obtained from the Bloomington *Drosophila* Stock Centre. The control strain used for all experiments was *w^1118^* (#60000, referred to as WT or +). The RNAi knockdown lines for CG14291 (v16897; referred to as IIIA here and as CG14291-IR1 in the original description of the model; (18)) and CG6903 (v109444; referred to as IIIC) were obtained from Vienna *Drosophila* Resource Centre (48).

Elav2-GAL4 and actin5c-GAL4 were used to drive the RNAi-mediated knockdown of CG14291 (*SGSH*) or CG6903 (*HGSNAT*) expression in neuronal and ubiquitous lines, respectively (18; L. Hewson Honours thesis). *Drosophila melanogaster* stocks were maintained in vials containing medium consisting of 1% (w/v) agar, 6% fresh yeast, 10% molasses, 1% glucose, 8.4% coarse semolina, 1.5% acid mix and 2.5% tegosept (referred to as control media). All flies were maintained at 18°C or 25°C with 60% humidity on a 12-hour light/dark cycle. They were turned into fresh vials with medium every 3-4 days. For drug treated flies, flies were raised on control media. After eclosion, adults were transferred into vials with the appropriate compound diluted in water at concentrations up to 100 mM mixed with control media. All assays were performed with male flies at 25°C. For larval experiments, flies were raised directly onto drug supplemented media.

### Quantitative PCR

Flies (ten pooled male flies at 0-3-days-post-eclosion) were homogenised using a glass mortar and pestle followed by aspiration with a 20G syringe in a total of 1 mL Trizol reagent (Thermo Fisher Scientific). Following centrifugation to remove cellular debris, total RNA was isolated and cDNA was generated using 1-μg of DNase I-treated RNA using a Superscript III first strand synthesis system kit and oligo(dT)20 primer (Thermo Fisher Scientific). Samples were then treated with RNase H (Invitrogen). Comparative gene expression of *sgsh/CG14291* or *hgsnat/CG6903* was determined on a QuantStudio 7 Flex (Thermo Fisher Scientific). Reactions were performed in triplicate in 384-well plates using Platinum SYBR Green qPCR SuperMix-UDG with ROX (Thermo Fisher Scientific) and primers (*CG14291* 5’-AGGCCAAGTACAATGGCACC-3’ / 5’-AGTCGAACAGTTGCTCGCG-3’; *CG6903* 5’-GAACACGCACTTCATGCTCCT-3’ / 5-CAGACCAGCGTGTTCCACG-3’ or rp49 comparator gene 5’-ATCGATATGCTAAGCTGTCGCAC-3’ / 5’-TGTCGATACCCTTGGGCTTG-3’) per the manufacturer’s instructions. Gene expression was calculated using the 2^ΔΔCt^ method and compared to a pooled WT control (either actin5c>+ and elav2>+).

### Tandem mass spectrometry

HS was quantitated using a modified method of He *et al*. (24). Briefly, five biological replicates each containing five larva or flies per genotype were homogenised in 500 μL 10% methanol using Lysing Matrix D tubes (MP Biomedicals, OH). Fly homogenates or cell extracts (10 μg total protein) were transferred to glass tubes and freeze-dried overnight using Rotational Vacuum Contractor RVC 2-33CD plus (Martin Christ) and Freeze-Dryer Alpha 2-4 LD plus (Martin Christ) at 37°C at 10 mbar and 352-377 x *g*. Following butanolysis at 100°C (HS) (49) and drying under nitrogen, the samples were reconstituted in 200 *μL* 100 ng/mL deuterated HS internal standard (*n*-butyl (2-amino-2-deoxy-α-D-glucopyranosyl)-(1→4)-(*n*-butyl-d_9_ α-D-glucopyranosid)uronate) (24) and centrifuged for 15 minutes at 16060x*g* on a Biofuge pico centrifuge (Heraeus, Germany). The supernatant was analysed using an API 4000 QTRAP mass spectrometer (AB/Sciex, Concord, Canada) in MRM mode measuring the transition *m/z* 468.245 to 162.077 (HS) and *m/z* 477.300 to 162.077 (internal standard) with online liquid chromatographic separation using an Acquity UPLC (Waters) (49). The peak areas were calculated using Analyst 1.6.2 (AB/Sciex). All samples were analysed in a random order interspersed every three injections with a blank injection of MilliQ water. All samples were quantified against a standard curve (1 and 2000 ng) of synthesised non-deuterated disaccharide *n*-butyl (2-amino-2-deoxy-α-D-glucopyranosyl)-(1→4)-(*n*-butyl α-D-glucopyranosid)uronate (24).

### *Drosophila* activity assays

The Drosophila Activity Monitoring system (TriKinetics, Inc) allows the recording of individual flies over a 1-week period for screening of altered behavioural profiles (50). An infrared beam is directed through the midpoint of each tube and the breaking of this beam is registered as an activity event. Briefly, flies were anaesthetised using carbon dioxide pads and placed individually in glass tubes containing 2% (w/v) agar and 6% (w/v) sucrose and then placed in the monitor. The flies were housed in a constant 12-hour light/12-hour dark cycle incubator (Digitherm) at 25°C. The FaasX software (M. Boudinot and F. Rouyer, Centre National de la Recherche Scientifique, Gif-sur-Yvette Cedex, France) was used to determine the average activity over the duration of the experiment.

### Climbing assays

Climbing assays were performed as previously described to characterise motor defects with the researcher blinded to the genotypes being tested (51). Assays were always carried out at the same time of day (12 pm) when male flies were transferred into a 500-mL measuring cylinder without anaesthetisation and the cylinder sealed with parafilm to prevent the flies from escaping. After 2 minutes of recovery, the flies were tapped to the bottom of measuring cylinder. After 25 seconds, the number of flies within each predefined area of the cylinder was counted. Flies that failed to climb above the 300-mL mark were considered to have a climbing defect. Assays were repeated in three consecutive trials for each cohort of flies, with a two-minute rest in between assays. The same cohort of flies was tested on days 1, 8 and 15.

### Social space assays

Social space assays were conducted as previously described (25, 52). Triangular chambers were custom-made with dimensions of 11.92 cm base, 14.5 cm height and 0.3 mm internal gap thickness (Gerald Scientific). Briefly, 40 male flies were transferred to a carbon dioxide pad for anaesthetisation and transferred into the chamber with horizontal orientation to account for the genotype differences in climbing ability and left for 20-30 minutes to settle. The social space chamber and a ruler for scale was photographed once the flies had settled. Digital images were imported in ImageJ software (NIH, rsbweb.nih.gov/ij) (53) and analysed for nearest neighbour distances (ImageJ plugin by YuXiong Mao). Genotypes were blinded while analysing the data. The social space assays were repeated in triplicate with different cohorts of the flies per genotype for each time-point.

### Lifespan assays

The lifespan of flies (n=50 males/genotype) was determined as previously described (26). Flies were transferred to fresh vials every 2-3 days and the number of deaths was recorded at each vial change. The maximum lifespan was calculated at the average age of the top 10% of flies from each cohort.

### Statistical analyses

All data are presented as mean ± SEM. Statistical analysis was completed using GraphPad Prism 7 (GraphPad Software, CA) or SPSS for Windows^®^ 15.0 (IBM Corporation, NY). Three-way ANOVA was used to determine statistical significance with genotype, drug and dose or genotype, drug and age as the independent factors. For climbing assays, repeated measures two-way ANOVA test with genotype and age or genotype and treatment as the independent factors were used. Log-rank (Mantel-Cox) tests were used for lifespan assays. Two-way ANOVA tests were used for other assays. The Bonferroni *post-hoc* test was applied to adjust for multiple comparisons between groups.

## Acknowledgements

This work was supported in part by the Sanfilippo Children’s Foundation (to VF, AL, MS, KH), the Science With Impact Fund (University of Queensland; to VF), and the Australian Research Council (DP170104431 to VF).

## Conflict of Interest Statement

The authors declare no conflict of interest.

---

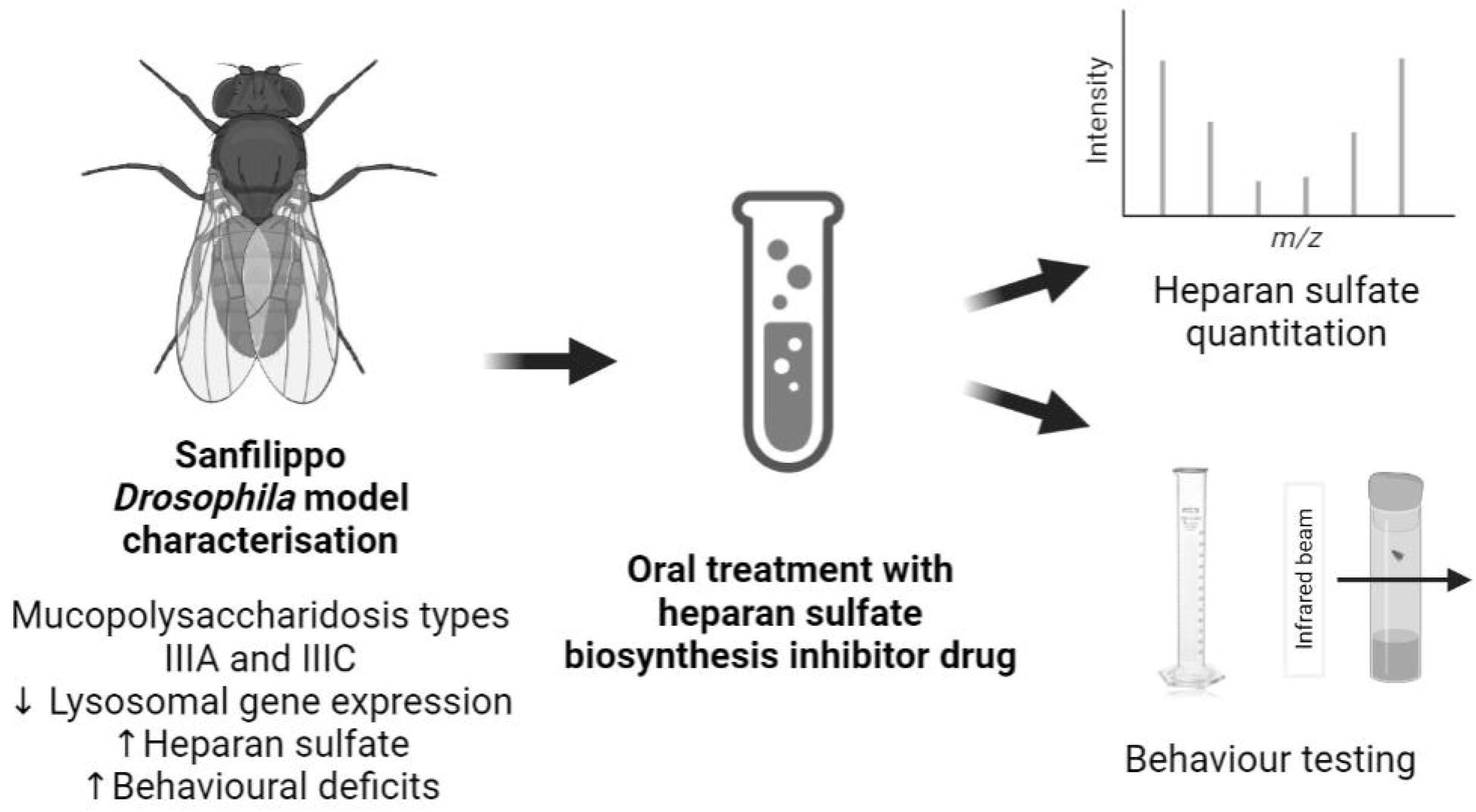

## Notes

### Competing Interest Statement

The authors have declared no competing interest.

