## Supplementary Information for "Substrate reduction therapy in a *Drosophila melanogaster* model of Sanfilippo syndrome"

### Table of contents

Supplementary Figure 1. Synthesis of 4-deoxy GlcNAc peracetate (compound 8) S3

Supplementary Figure 2. Phenotypic screening in Sanfilippo *Drosophil*a. S4

Supplementary Figure 3. Effect of drug concentration of life span. S5

Supplementary Figure 4. Copies of $^{13}$C NMR and/or $^{1}$H and spectra for compounds **2-8** S6-S19

Supplementary Methods. Synthesis of 4-deoxy-GlcNAc peracetate S20-S25

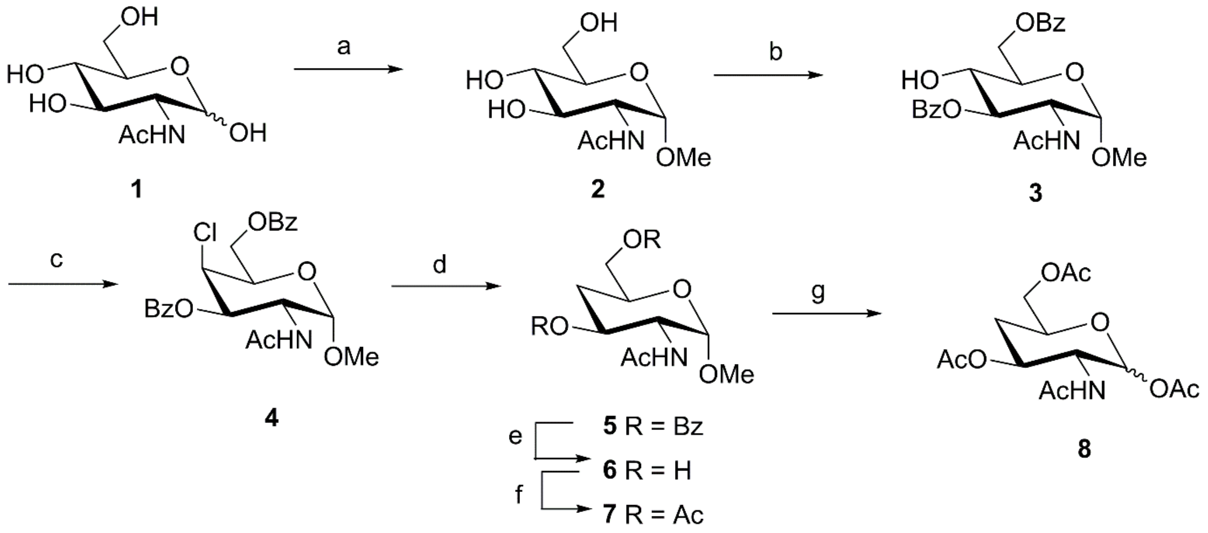


**Supplementary Figure 1. Scheme 1 - Synthesis of 4-deoxy GlcNAc peracetate (compound 8)**. *Reagents and conditions*: a) MeOH, Amberlite® IR120 ($H^+$), reflux, 24 h, 98%; b) BzCl, pyridine, -40 ºC, 67%; c) $SO_2Cl_2$, pyridine, 0 ºC, 3h, 100%; d) $Bu_3SnH$, VASO, toluene, reflux 24 h, 98%; e) KOH, MeOH, r.t., 24h, 90%; f) $Ac_2O$, pyridine, DMAP, r.t., 24 h, 100%; g) $Ac_2O$, conc.$H_2SO_4$, r.t., 4 h, 91%.

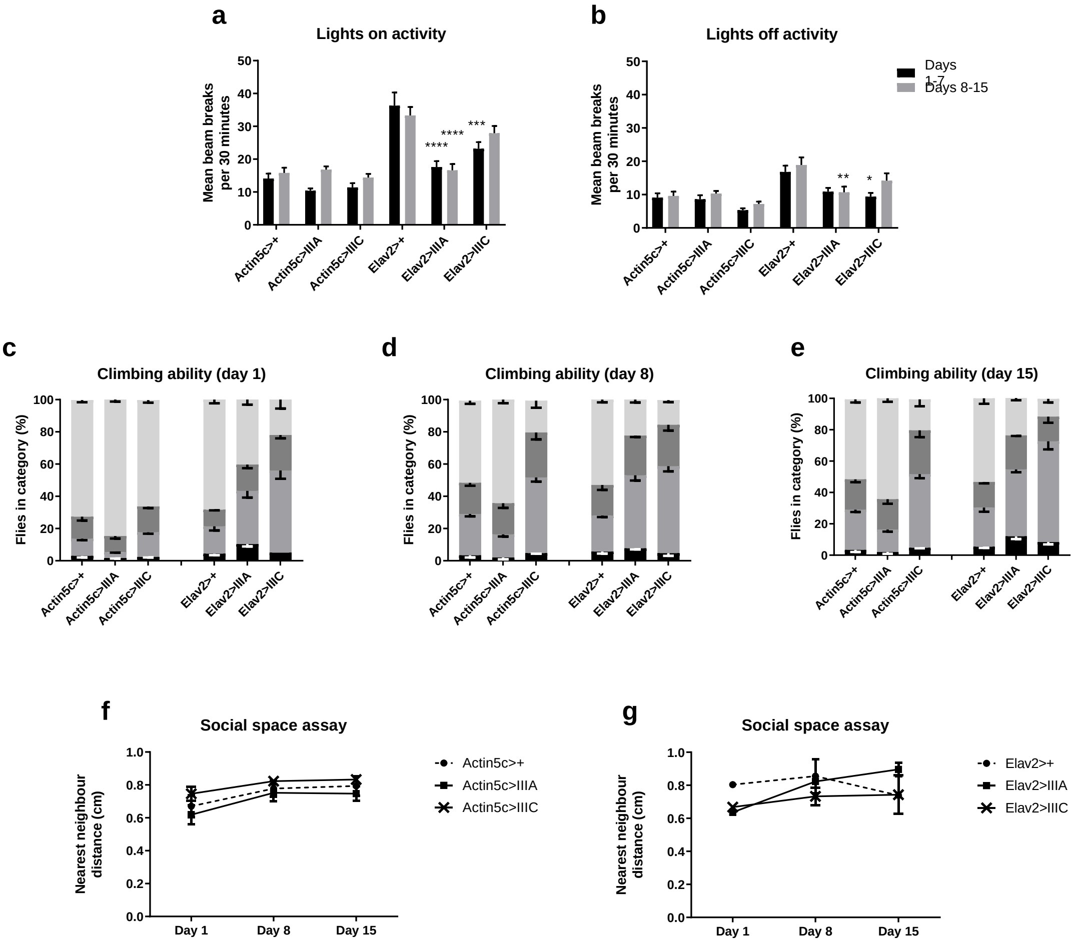


**Supplementary Figure 2. Phenotypic screening in Sanfilippo *Drosophil*a.** (a, b) Individual male flies were housed in a *Drosophila* Activity Monitoring system over a 7-day period. The average number of beam breaks was used as an measure of locomotor activity (n=14-16 males/genotype/time-point). Flies were either 1- or 8-days post-eclosion on the first day of recording. (c-e) At 1-, 8- and 15-days post-eclosion, flies were transferred to a 500 ml measuring cylinder, tapped to the bottom, and then allowed to climb for 25-seconds. The number of flies in each region was counted (<50 ml, 50–300 ml, 300–500 ml and >500 ml; n=110-179 male flies/genotype/time-point). A failure to climb is indicative of a motor defect. (f, g) For each test, flies were briefly anaesthetised and transferred into triangular chambers (n=40). After an acclimatisation period, the position of the flies was digitally imaged and the distance from one fly to its nearest neighbour was calculated using Image J. Each social space assay was conducted in triplicate.

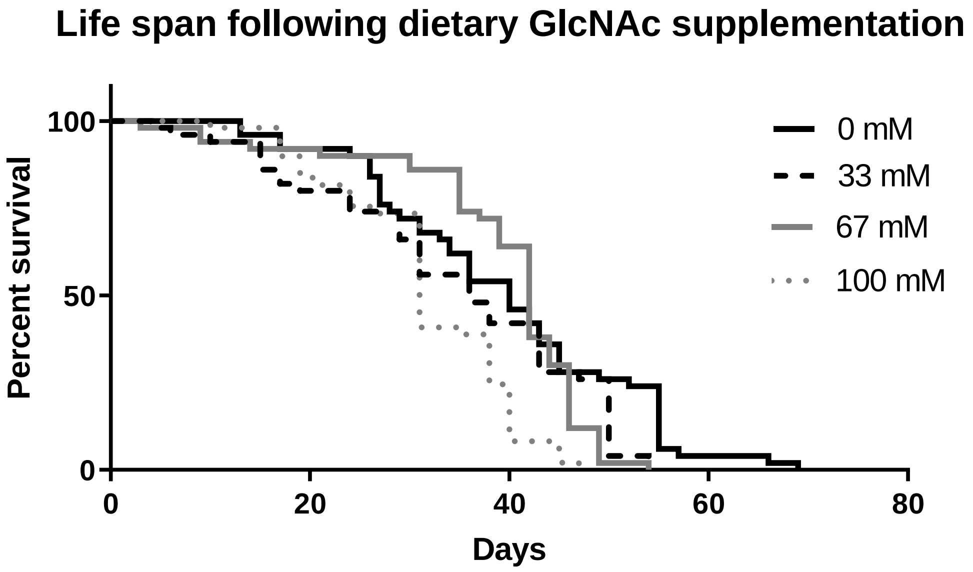


**Supplementary Figure 3. Effect of drug concentration of life span.** To examine drug toxicity, GlcNAc was orally delivered to actin5c>+ flies in food agar at one of four doses. Flies were transferred to fresh vials every 2-3 days and the number of deaths was recorded at each food change. Flies fed 100 mM GlcNAc had a significant decrease in life span compared to compared to the vehicle-supplemented controls.

**Supplementary Figure 4.** $^{13}$C NMR and/or $^{1}$H and spectra for compounds **2-8.**

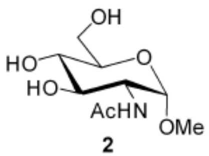

**2**

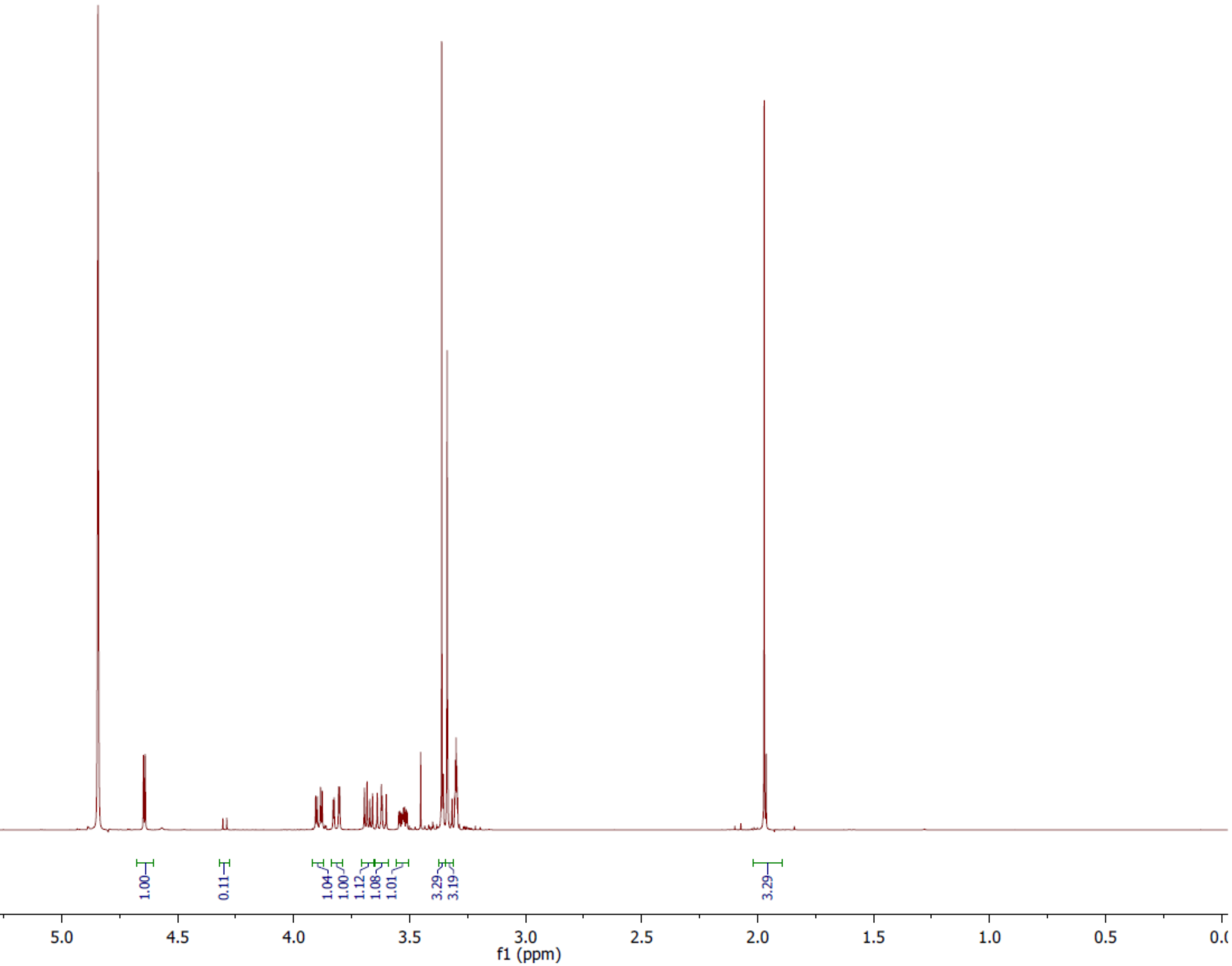

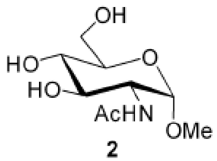

**2**

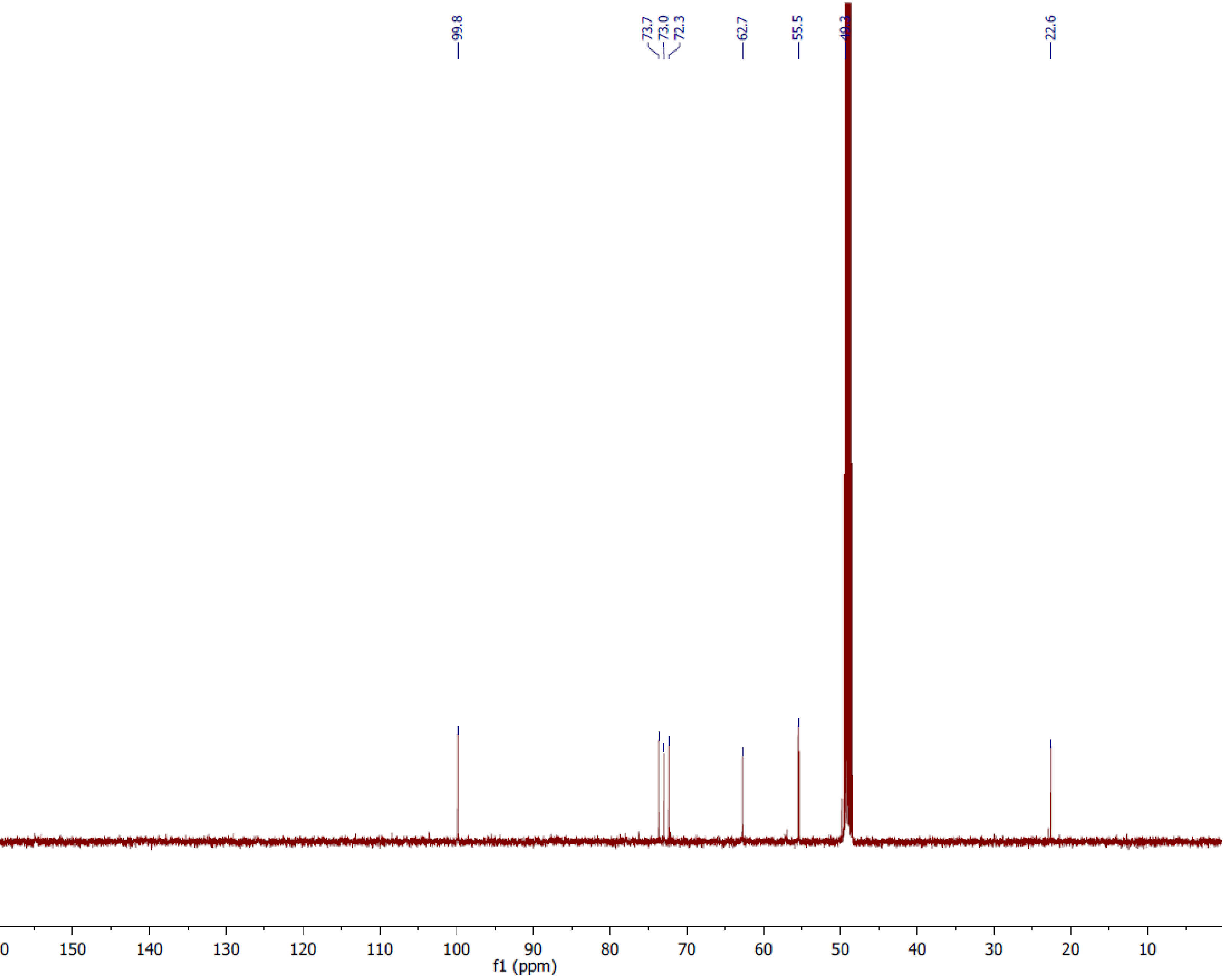
99.8
73.7
73.0
72.3
62.7
55.5
49.3
22.6
180
170
160
150
140
130
120
110
100
90
80
70
60
50
40
30
20
10
f1 (ppm)

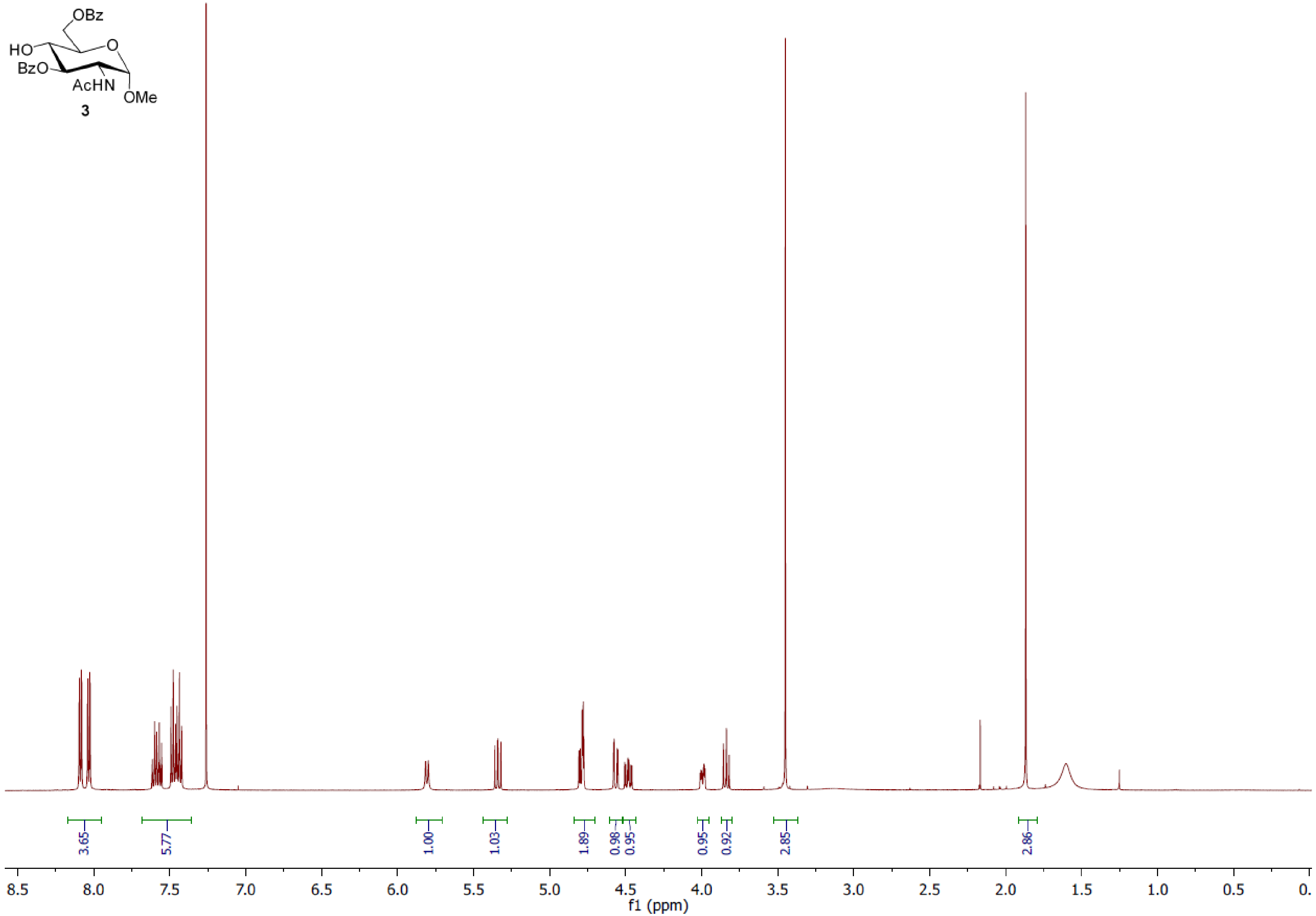
OBz
HO
O
BzO
AcHN
OMe
3
3.65
5.77
1.00
1.03
1.89
0.98
0.95
0.95
0.92
2.85
2.86
8.5
8.0
7.5
7.0
6.5
6.0
5.5
5.0
4.5
4.0
3.5
3.0
2.5
2.0
1.5
1.0
0.5
0.
f1 (ppm)

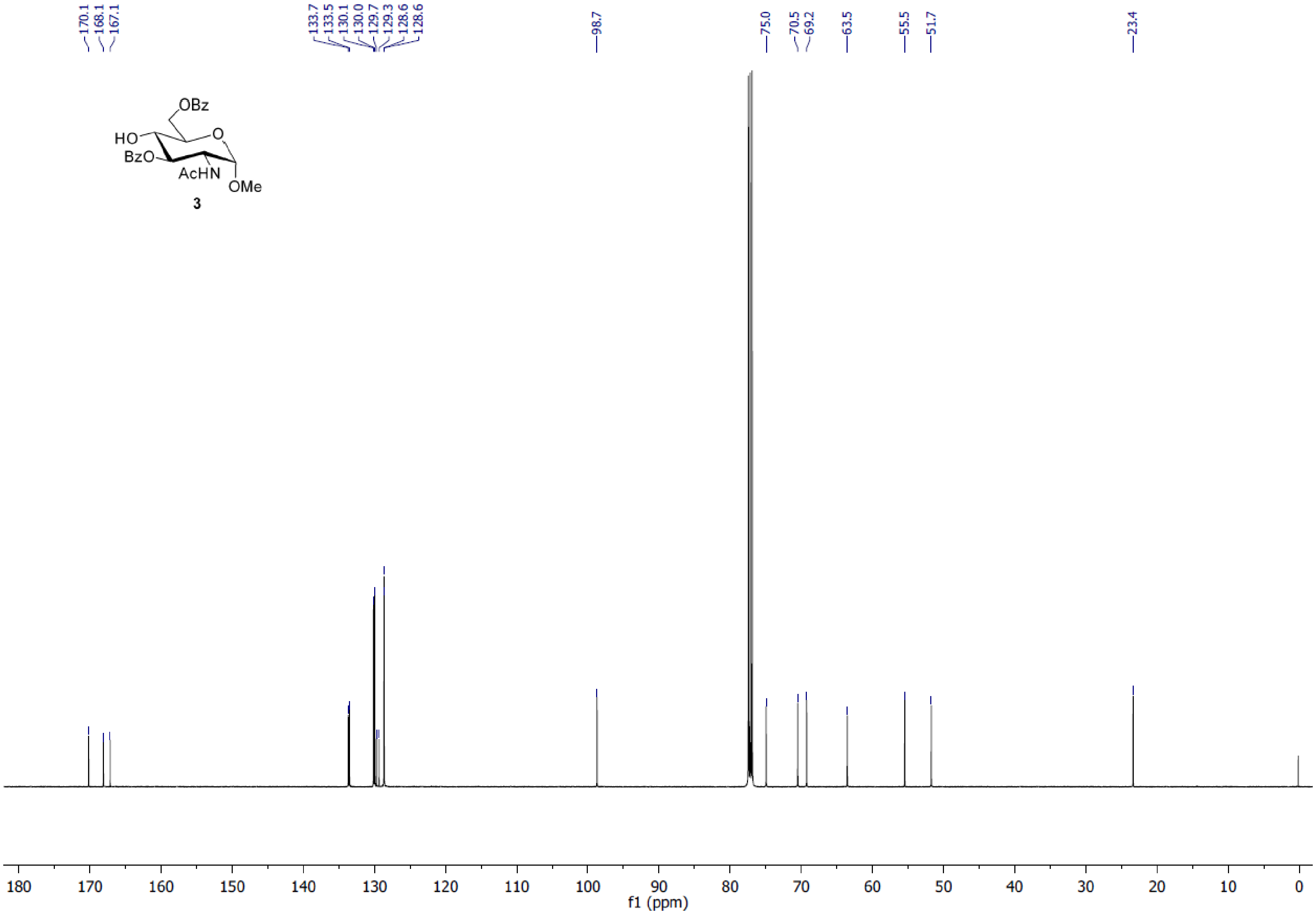
170.1
168.1
167.1
133.7
133.5
130.1
130.0
129.7
129.3
128.6
128.6
98.7
75.0
70.5
69.2
63.5
55.5
51.7
23.4
OBz
HO
BzO
O
AcHN
OMe
3
180 170 160 150 140 130 120 110 100 90 80 70 60 50 40 30 20 10 0
f1 (ppm)

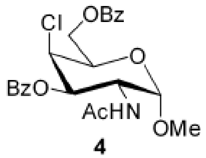
Cl
OBz
O
BzO
AcHN
OMe

**4**

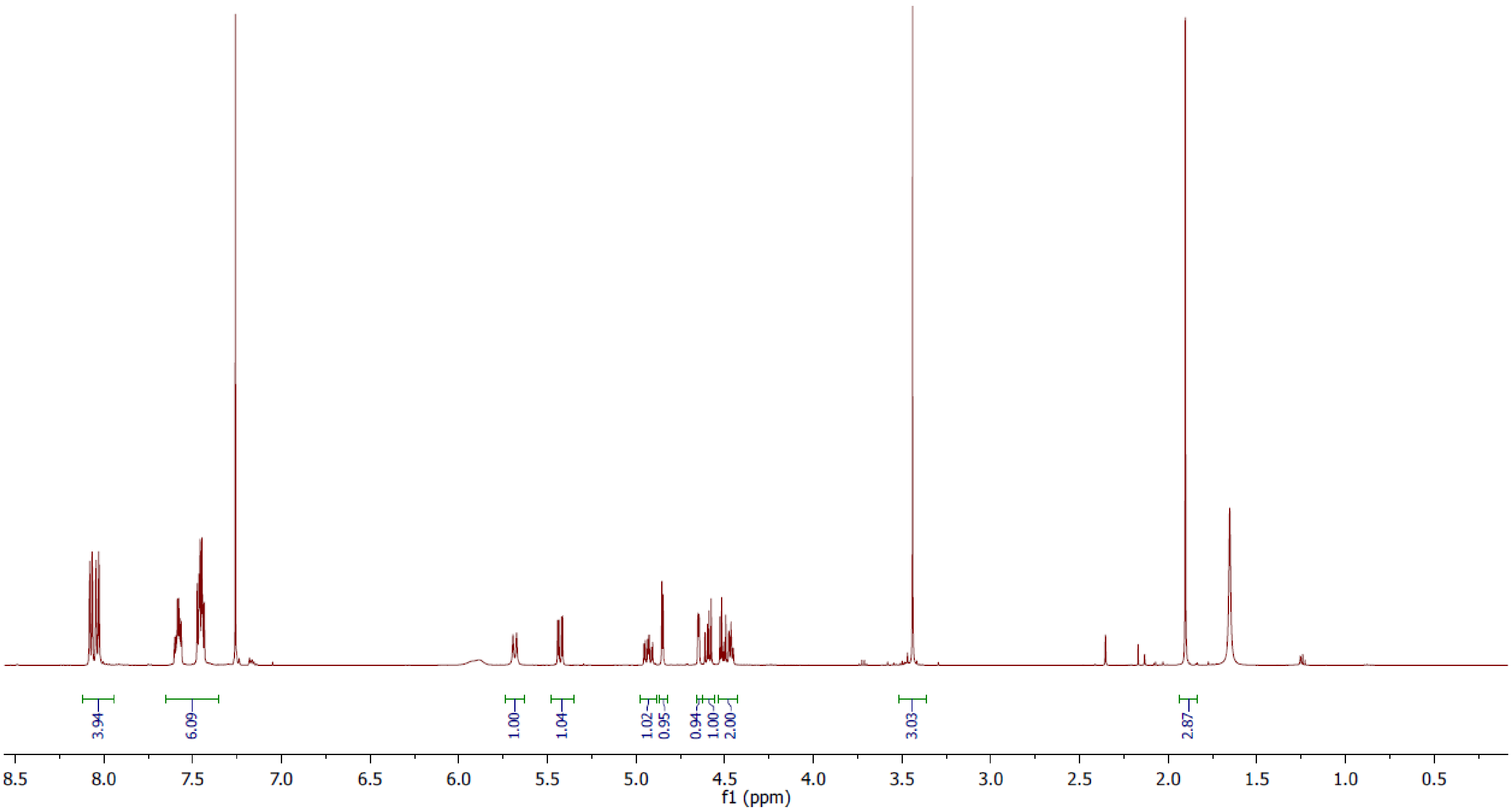
3.94
6.09
1.00
1.04
1.02
0.95
0.94
1.00
2.00
3.03
2.87
8.5
8.0
7.5
7.0
6.5
6.0
5.5
5.0
4.5
4.0
3.5
3.0
2.5
2.0
1.5
1.0
0.5
f1 (ppm)

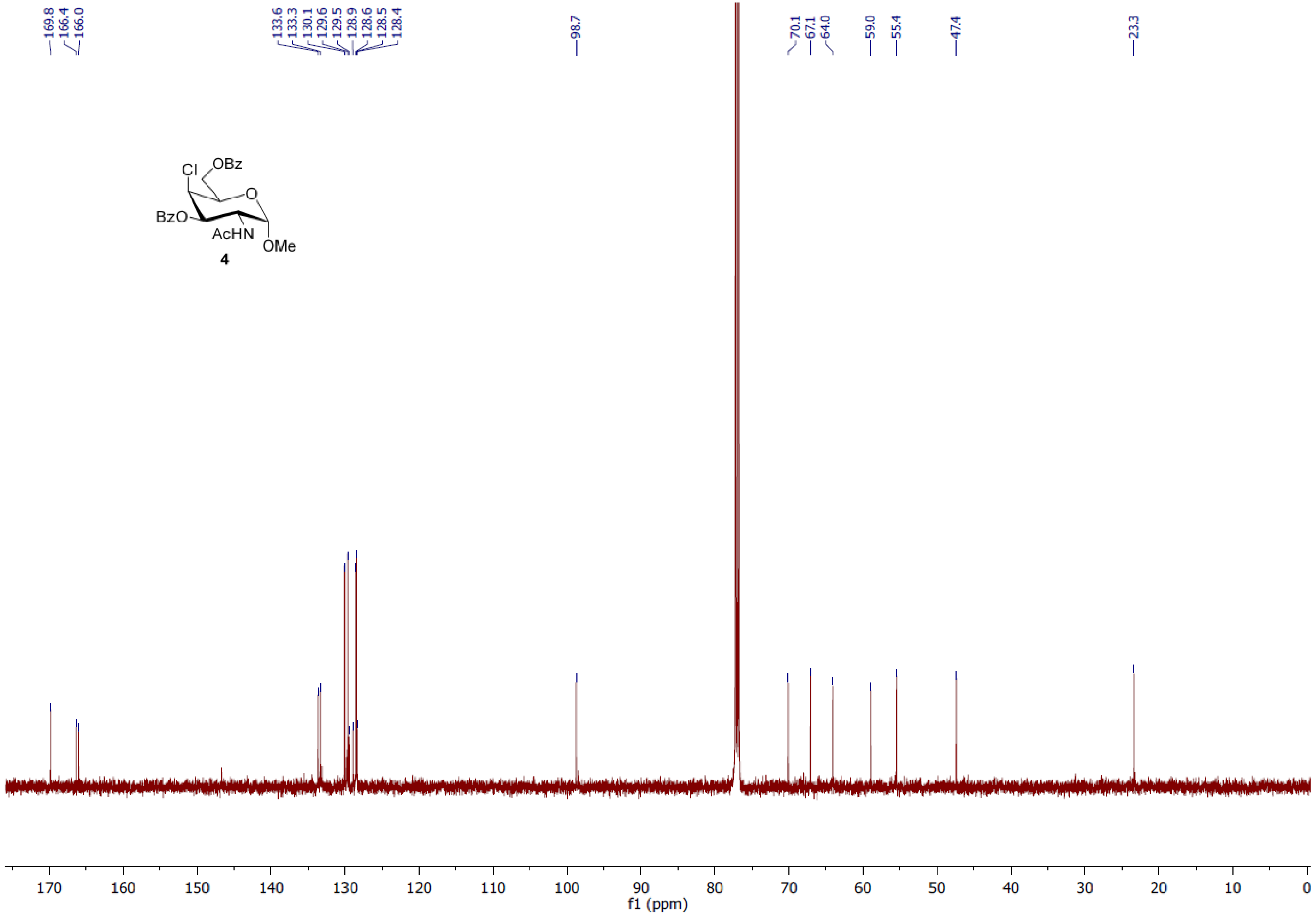
169.8
166.4
166.0
133.6
133.3
130.1
129.6
129.5
128.9
128.6
128.5
128.4
98.7
70.1
67.1
64.0
59.0
55.4
47.4
23.3
Cl
OBz
O
BzO
AcHN
OMe
4
170
160
150
140
130
120
110
100
90
80
70
60
50
40
30
20
10
0
f1 (ppm)

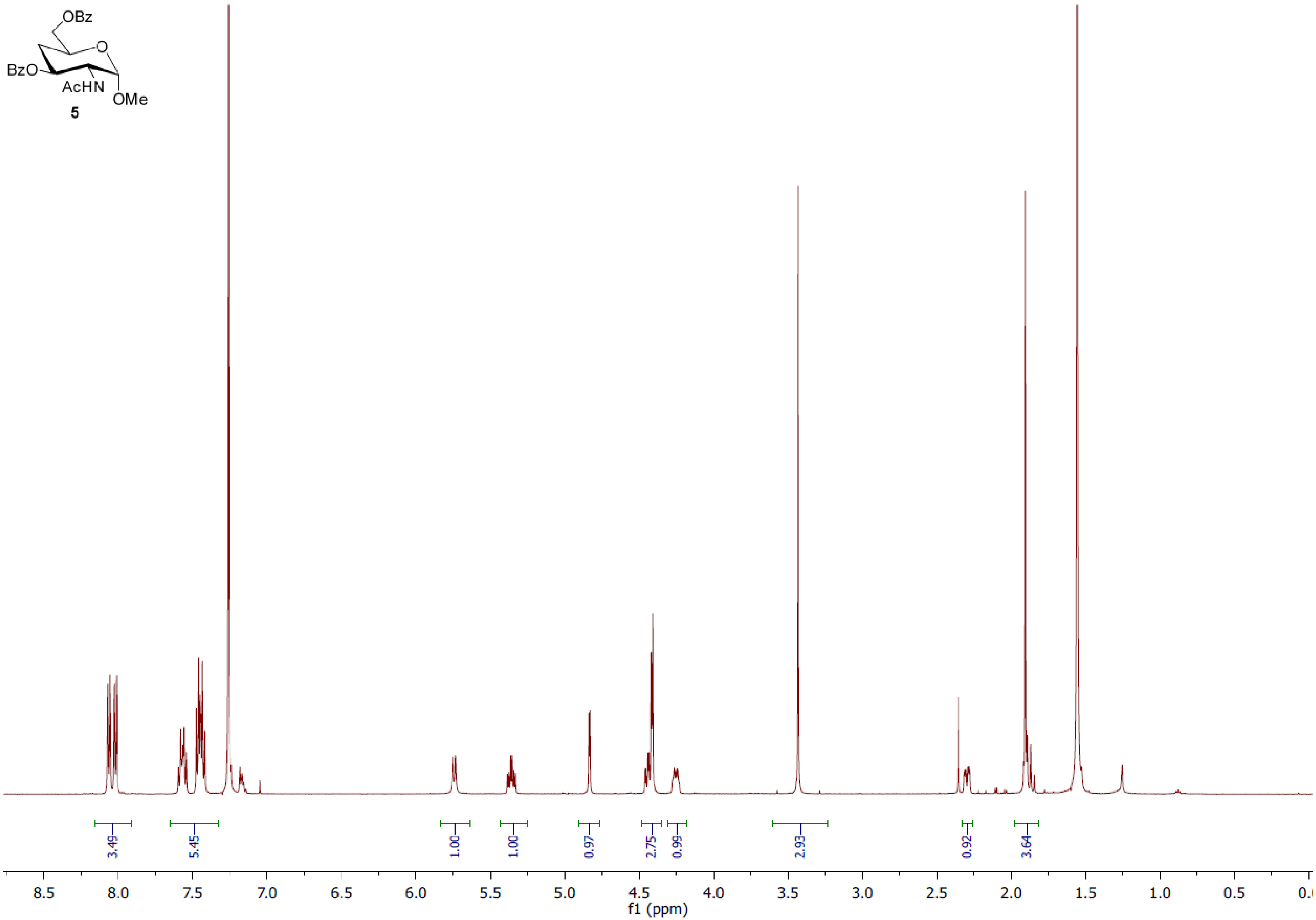
OBz
BzO
O
AcHN
OMe
5
3.49
5.45
1.00
1.00
0.97
2.75
0.99
2.93
0.92
3.64
8.5
8.0
7.5
7.0
6.5
6.0
5.5
5.0
4.5
4.0
3.5
3.0
2.5
2.0
1.5
1.0
0.5
f1 (ppm)

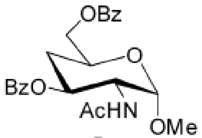
OBz
O
BzO
AcHN
OMe

**5**

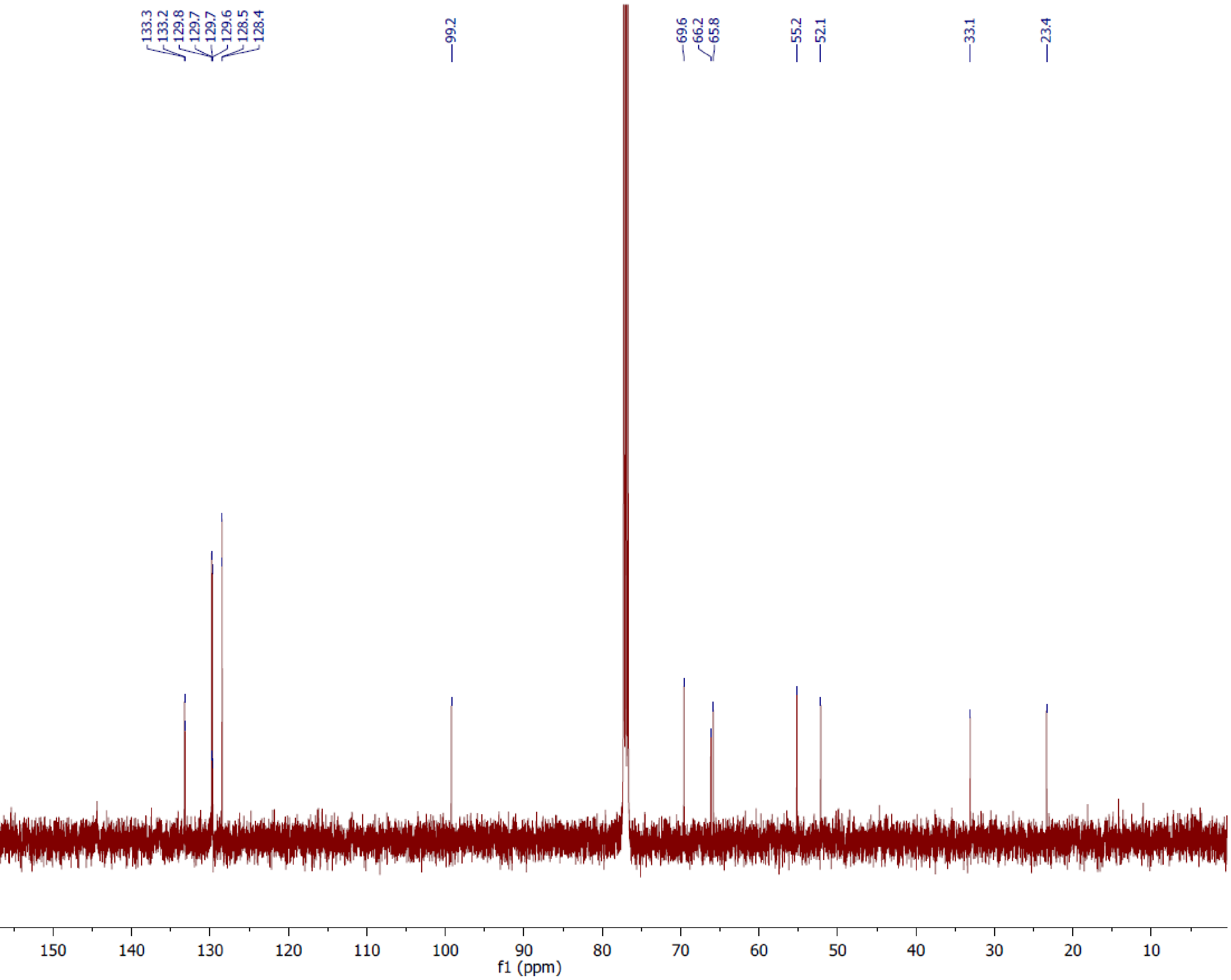
133.3
133.2
129.8
129.7
129.7
129.6
128.5
128.4
99.2
69.6
66.2
65.8
55.2
52.1
33.1
23.4
170
160
150
140
130
120
110
100
90
80
70
60
50
40
30
20
10
f1 (ppm)

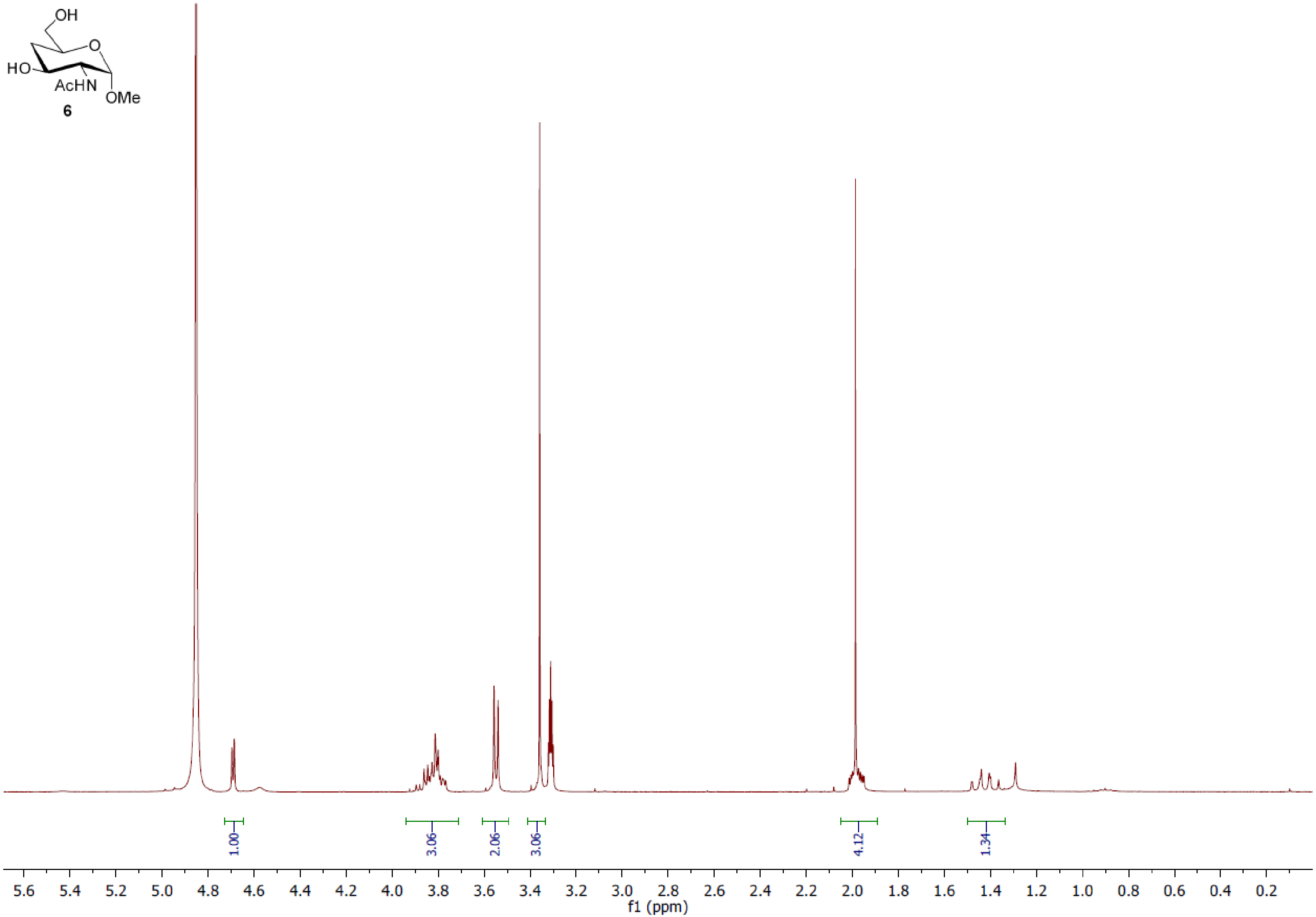
OH
O
HO
AcHN
OMe
6
1.00
3.06
2.06
3.06
4.12
1.34
5.6 5.4 5.2 5.0 4.8 4.6 4.4 4.2 4.0 3.8 3.6 3.4 3.2 3.0 2.8 2.6 2.4 2.2 2.0 1.8 1.6 1.4 1.2 1.0 0.8 0.6 0.4 0.2
f1 (ppm)

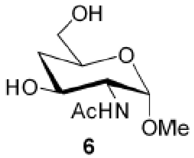
OH
O
HO
AcHN
OMe

**6**

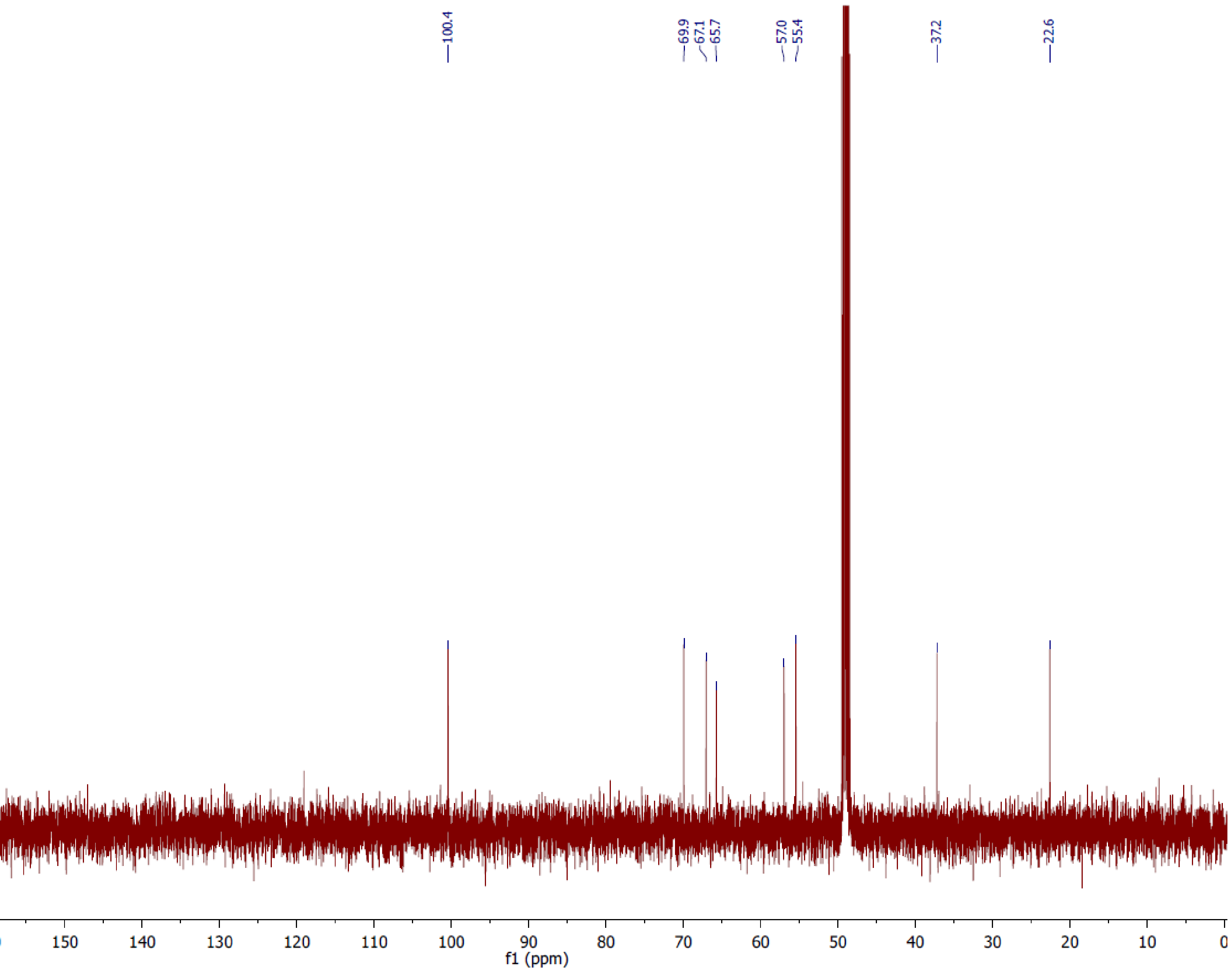
100.4
69.9
67.1
65.7
57.0
55.4
37.2
22.6
170
160
150
140
130
120
110
100
90
80
70
60
50
40
30
20
10
0
f1 (ppm)

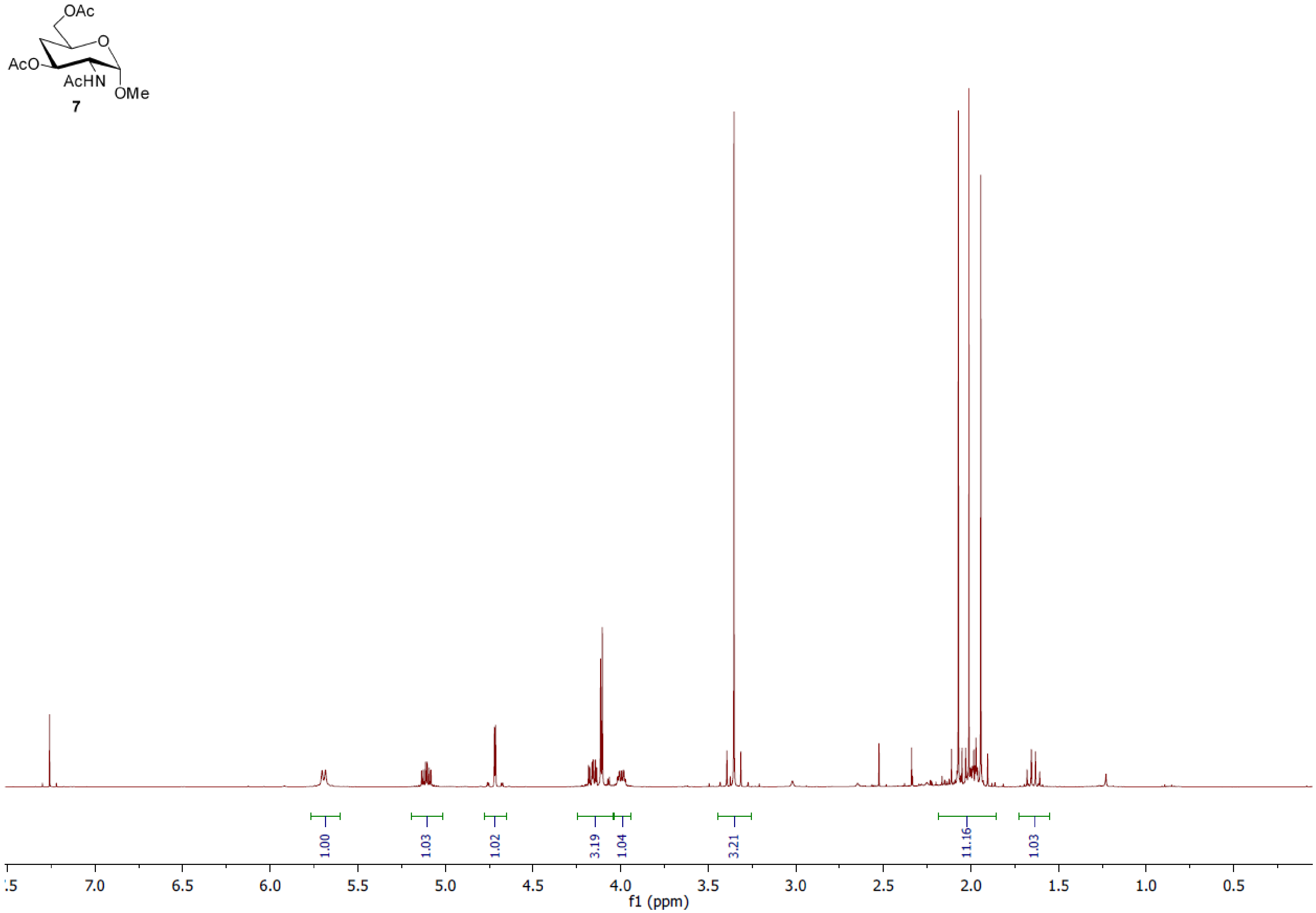
OAc
O
AcO
AcHN
OMe
7
1.00
1.03
1.02
3.19
1.04
3.21
11.16
1.03
7.0
6.5
6.0
5.5
5.0
4.5
4.0
3.5
3.0
2.5
2.0
1.5
1.0
0.5
f1 (ppm)

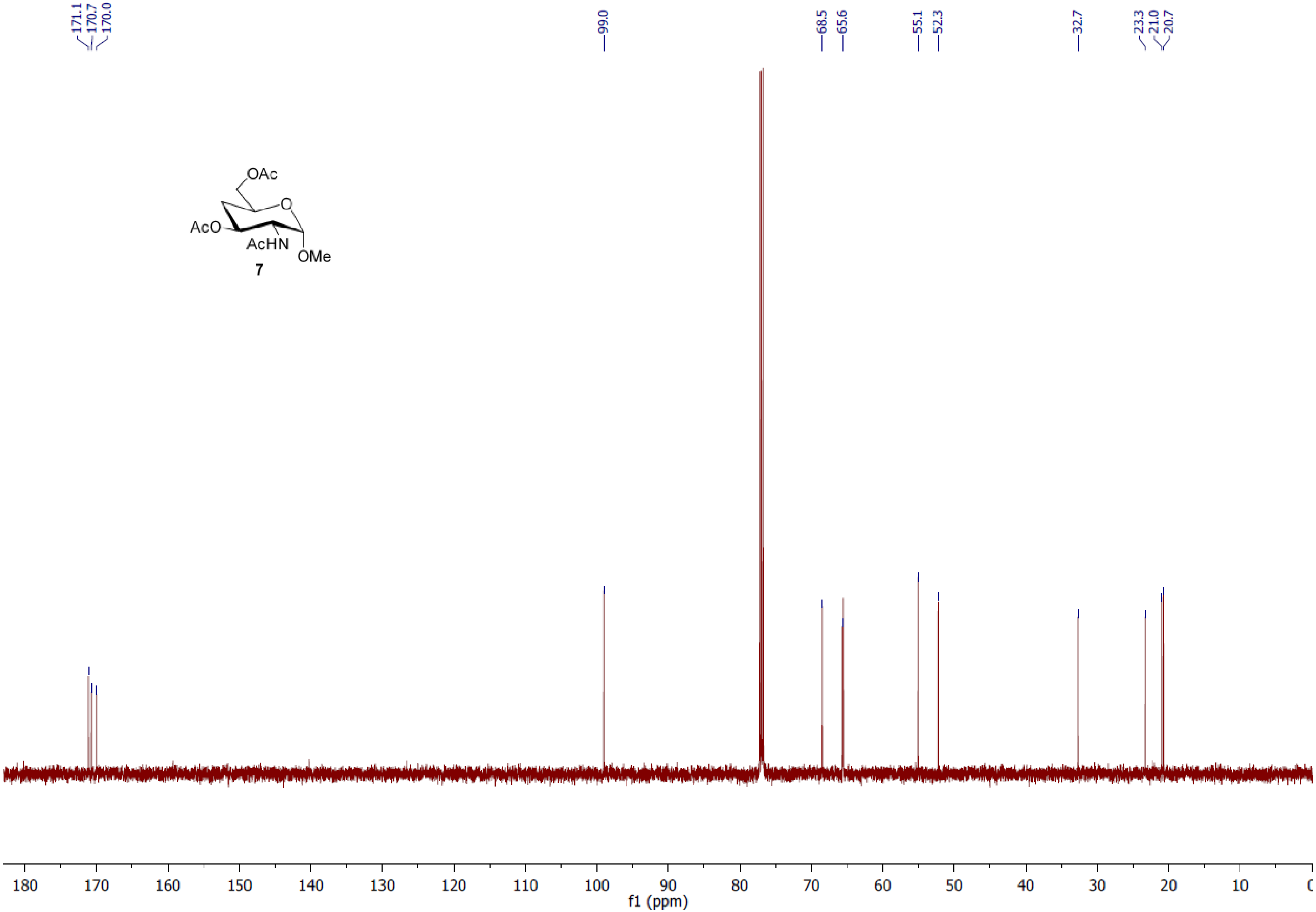
171.1
170.7
170.0
99.0
68.5
65.6
55.1
52.3
32.7
23.3
21.0
20.7
OAc
AcO
AcHN
OMe
O
7
f1 (ppm)

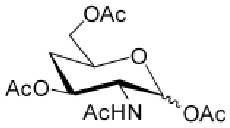
OAc
O
AcO
AcHN
OAc

**8**

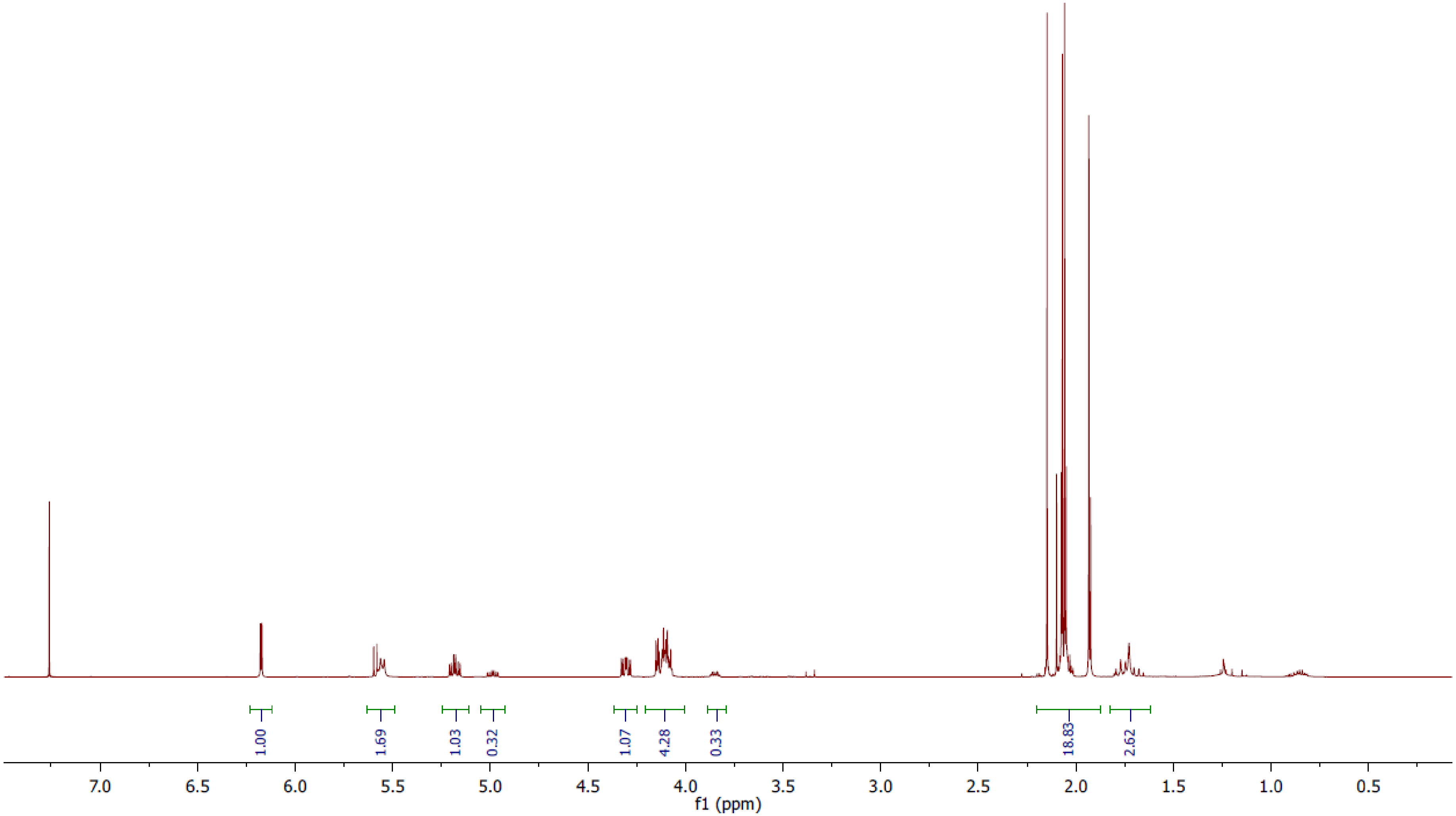
1.00
1.69
1.03
0.32
1.07
4.28
0.33
18.83
2.62
7.0
6.5
6.0
5.5
5.0
4.5
4.0
3.5
3.0
2.5
2.0
1.5
1.0
0.5
f1 (ppm)

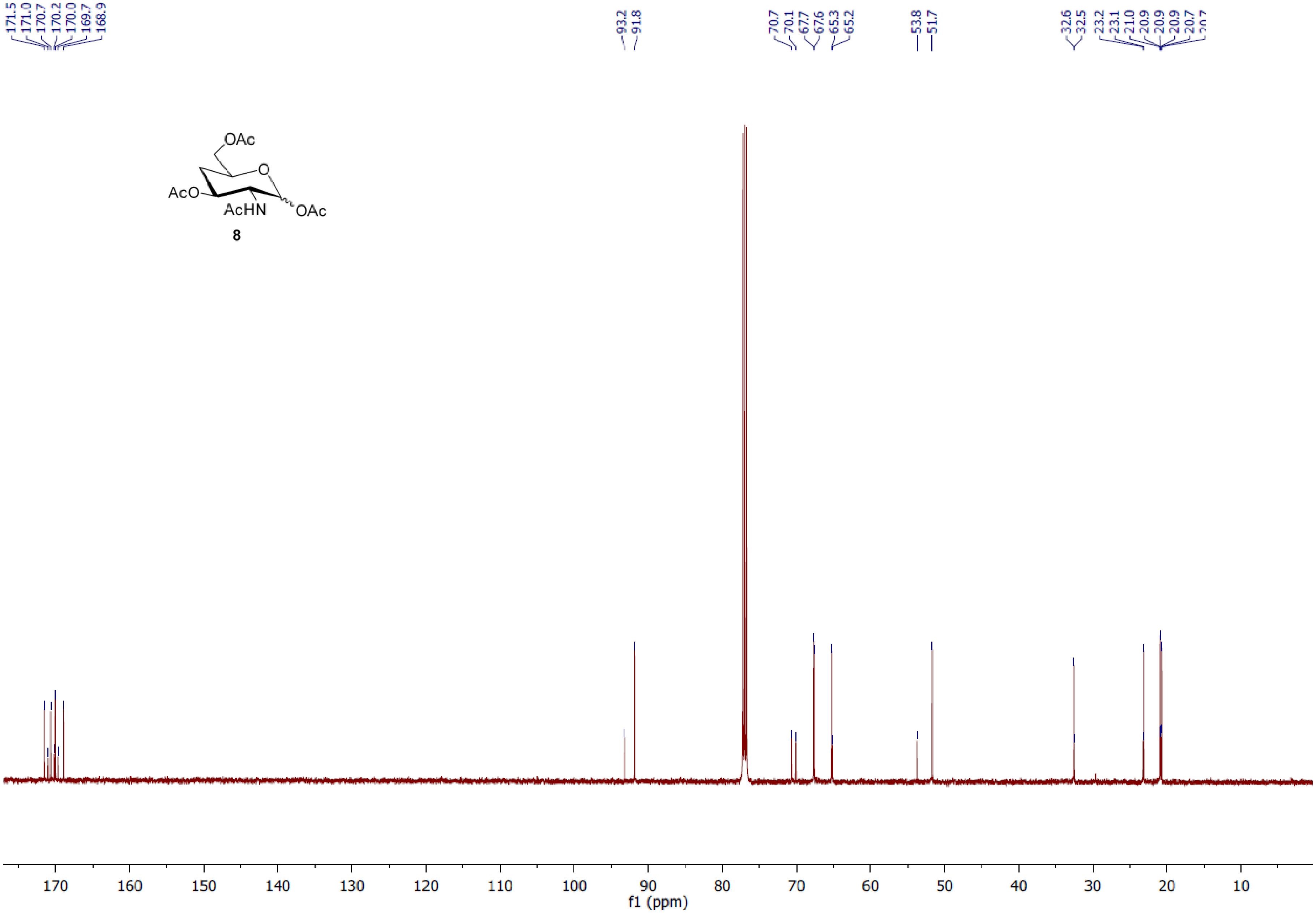

8

171.5
171.0
170.7
170.2
170.0
169.7
168.9
93.2
91.8
70.7
70.1
67.7
67.6
65.3
65.2
53.8
51.7
32.6
32.5
23.2
23.1
21.0
20.9
20.9
20.9
20.7
170 160 150 140 130 120 110 100 90 80 70 60 50 40 30 20 10
f1 (ppm)

#### Supplementary Methods - Synthesis

##### General methods

Melting points were determined on a DigiMelt MSRS apparatus. Optical rotations were determined on a JASCO P-2000 polarimeter at ambient temperature and are given in units of $10^{-1}$ deg $cm^2$ $g^{-1}$. $^1H$ and $^{13}C$ NMR spectra were recorded on a Bruker Avance 500 MHz spectrometer at 20 °C. The residual solvent peaks ($CDCl_3$: $\delta_H$ 7.24 and $\delta_C$ 77.0; $CD_3OD$: $\delta_H$ 4.78 and $\delta_C$ 49.0) served as internal standards. Coupling constants in Hz were measured from one-dimensional spectra. Spectral assignments were aided by 2D (COSY, HSQC) experiments. Low resolution mass spectra were acquired on a Bruker HCT 3D mass spectrometer. All chemicals were purchased as reagent grade and used without further purification. Solvents were distilled before use. Reactions were monitored by analytical thin layer chromatography (TLC) on silica gel 60 $F_{254}$ plates and visualized by charring with 10% sulfuric acid in ethanol or by treatment with *p*-anisaldehyde stain.

##### Methyl 2-acetamido-2-deoxy-α-D-glucopyranoside (2)

Amberlite IR120 ($H^+$) resin was added to a solution of *N*-acetyl-D-glucosamine **1** (10.32 g, 46.64 mmol) in anhydrous MeOH (220 mL). The mixture was heated at reflux for 24 h and then cooled to r.t. The resin was removed by filtration and washed with MeOH (5 × 35 mL). The filtrate and washings were evaporated to dryness and co-evaporated with toluene under reduced pressure to give the glycoside **2** as a white powder (10.82 g, 98%). $R_f$ = 0.45 (3:1 EtOAc/MeOH), m.p. 187 – 189 °C (lit.[1] 187 – 188 °C). The ratio of α:β anomers was 10:1 by $^1H$ NMR spectroscopy. **2α**: $^1H$ NMR (500 MHz, $CD_3OD$): δ 4.64 (d, 1H, $J_{1,2}$ = 3.6 Hz, H-1), 3.89 (dd, 1H, $J_{2,3}$ = 10.7 Hz, H-2), 3.81 (dd, A part of ABX, 1H, $J_{5,6a}$ = 2.4, $J_{6a,6b}$ = 11.9 Hz, H-6a), 3.68 (dd, B part of ABX, 1H, $J_{5,6b}$ = 5.8 Hz, H-6b), 3.62 (dd, 1H, $J_{3,4}$ = 8.8 Hz, H-3), 3.53 (ddd, 1H, $J_{4,5}$ = 9.9 Hz, H-5), 3.36 (s, 3H, OMe), 3.34 (dd, 1H, H-4), 1.97 (s, 3H, NHCOC$H_3$); $^{13}C$ NMR (125 MHz, $CD_3OD$): δ = 173.7 (C=O), 99.8 (C-1), 73.7 (C-5), 73.0 (C-3), 72.3 (C-4), 62.7 (C-6), 55.5 (C-2), 55.4 (OMe), 22.6 (NHCO*C*$H_3$).

**Methyl 2-acetamido-3,6-di-*O*-benzoyl-2-deoxy-α-D-glucopyranoside (3)**

The triol **2** (2.04 g, 8.7 mmol) was dissolved in anhydrous pyridine (20 mL) and cooled to – 40 °C. Benzoyl chloride (2.1 mL, 18.2 mmol, 2.1 eq.) was added dropwise over two min and the reaction mixture was stirred at – 40 °C for 1.5 h. MeOH (5 mL) was then added and the mixture was concentrated and co-evaporated with toluene. The residue was purified by flash chromatography (EtOAc/hexane, 2:1) to give the dibenzoate **3** as a white solid (2.58 g, 67%), m.p. 147 – 148 °C (lit.[2] 147-149 °C [EtOAc-pet. ether]); $[\alpha]_D$ +104.6 (*c* 1.7, $CHCl_3$; lit.[2] +110.4); $R_f$ = 0.35 (EtOAc/hexane, 2:1). $^1$H NMR (500 MHz, $CDCl_3$): δ 8.09 – 8.02 (m, 4H, Ph), 7.61 – 7.55 (m, 2H, Ph), 7.49 – 7.42 (m, 4H, Ph), 5.81 (d, 1H, $J_{NH,2}$ = 9.6 Hz, NH), 5.34 (dd, 1H, $J_{2,3}$ = 9.1 Hz, $J_{3,4}$ = 9.7 Hz, H-3), 4.79 (dd, A part of ABX, 1H, $J_{5,6a}$ = 4.2 Hz, $J_{6a,6b}$ = 12.2 Hz, H-6a), 4.78 (d, 1H, $J_{1,2}$ = 3.5 Hz, H-1), 4.56 (dd, 1H, $J_{5,6b}$ = 2.3 Hz, H-6b), 4.48 (ddd, 1H, $J_{NH,2}$ = 9.6 Hz, H-2), 4.00 (ddd, 1H, $J_{4,5}$ = 9.7 Hz, H-5), 3.84 (dd, 1H, H-4), 3.45 (s, 3H, OMe), 1.87 (s, 3H, NHCOC*H*$_3$); $^{13}$C NMR (125 MHz, $CDCl_3$): δ 170.1, 168.1, 167.1 (3 × C=O), 133.7, 133.5, 130.1, 130.0, 129.7, 129.3, 128.7, 128.6 (Ph), 98.7 (C-1), 75.0 (C-3), 70.5 (C-5), 69.2 (C-4), 63.5 (C-6), 55.5 (OMe), 51.7 (C-2), 23.4 (NHCO*C*H$_3$). LRMS: *m/z* = 443.96 $[M+H]^+$, 466.20 $[M+Na]^+$.

**Methyl 2-acetamido-3,6-di-*O*-benzoyl-4-chloro-2,4-dideoxy-α-D-galactopyranoside (4)**

To a solution of the alcohol **3** (224 mg, 0.50 mmol) in anhydrous pyridine (2.5 mL) at 0 °C under Ar was added $SO_2Cl_2$ (53 μL, 0.66 mmol). This solution was stirred at 0 °C for 3 h, then at r.t. for 1.5 h. The solvent was evaporated under reduced pressure, and the residue was dissolved in $CHCl_3$, and washed with 2 M HCl (3 × 5 mL). The organic layer was dried ($MgSO_4$), filtered and concentrated to give the chloride **4** as light yellow solid (239 mg, 100%), used without further purification in the next step. $R_f$ = 0.31 (EtOAc/ *n*-hexane, 2:1); m.p. 178 – 184 °C; $^1$H NMR (500 MHz, $CDCl_3$): δ 8.08 – 8.03 (m, 4H, Ph), 7.60 – 7.56 (m, 2H, Ph), 7.47 – 7.43 (m, 4H, Ph), 5.65 (d, 1H, $J_{2,NH}$ = 9.7 Hz, NH), 5.43 (dd, 1H, $J_{2,3}$ = 10.9 Hz, $J_{3,4}$ = 3.6 Hz, H-3), 4.93 (ddd, 1H, $J_{1,2}$ = 3.6 Hz, H-2), 4.85 (d, 1H, H-1), 4.65 (dd, 1H, $J_{4,5}$ = 1.3 Hz, H-4), 4.60 (dd, A part of ABX, 1H, $J_{5,6a}$ = 6.9 Hz, $J_{6a,6b}$ = 11.3 Hz,

H-6a), 4.51 (dd, B part of ABX, 1H, $J_{5,6b}$ = 5.4 Hz, H-6b), 4.46 (ddd, 1H, H-5), 3.44 (s, 3H, OMe), 1.90 (s, 3H, NHCOC$H_3$); $^{13}$C NMR (125 MHz, $CDCl_3$): δ 169.8, 166.4, 166.0 (3 × C=O), 133.6, 133.3, 130.1, 129.7, 129.6, 129.5, 128.9, 128.6, 128.4 (Ph), 98.7 (C-1), 70.1 (C-3), 67.1 (C-5), 64.0 (C-6), 58.9 (C-4), 55.4 (OMe), 47.3 (C-2), 23.3 (NHCO*C*$H_3$). The NMR data were in accord with the literature.[3] LRMS: *m/z* = 461.96 $[M+H]^+$, 484.12 $[M+Na]^+$.

**Methyl 2-acetamido-3,6-di-*O*-benzoyl-2,4-dideoxy-α-D-*xylo*-hexopyranoside (5)**

A solution of the chloride **4** (3.05 g, 6.61 mmol), $Bu_3SnH$ (3.4 mL, 12.6 mmol) and catalytic amount of VASO (1,1'-azobis(cyclohexanecarbonitrile), 22 mg) in anhydrous toluene (30 mL) under Ar was heated at reflux for 24 h. The solution was evaporated and the residue partitioned between acetonitrile and *n*-hexane. Concentration of the acetonitrile phase followed by purification of the residue by flash chromatography (EtOAc/ *n*-hexane, 1:1) gave the 4-deoxy hexoside **5** as a white solid (2.81 g, 98%), m.p. 160–161 °C (lit.[2] 166–167 °C [EtOAc]); $[\alpha]_D$ +95.4 (*c* 0.74, $CHCl_3$; lit.[2] +106.8); $R_f$ = 0.27 (EtOAc/ *n*-hexane, 1: 1); TLCs for this reaction were visualized with *p*-anisaldehyde stain (**4** and **5** had the same $R_f$ but stained a different colour). $^1$H NMR (500 MHz, $CDCl_3$): δ 8.06 – 8.00 (m, 4H, Ph), 7.59 – 7.54 (m, 2H, Ph), 7.47 – 7.42 (m, 4H, Ph), 5.73 (d, 1H, $J_{NH,2}$ = 9.5 Hz, NH), 5.35 (ddd, 1H, $J_{3,4eq}$ = 4.9 Hz, $J_{3,4ax}$ = $J_{2,3}$ = 11 Hz, H-3), 4.83 (d, 1H, $J_{1,2}$ = 3.6 Hz, H-1), 4.47-4.40 (m, 3H, H-2, H-6a, 6b), 4.27 – 4.23 (m, 1H, H-5), 3.43 (s, 3H, OMe), 2.30 (ddd, 1H, $J_{4eq,5}$ = 2.0 Hz, $J_{4ax,4eq}$ = 12.5 Hz, H-4eq), 1.90 (s, 3H, Ac), 1.92-1.84 (m, 1H, H-4ax); $^{13}$C NMR (125 MHz, $CDCl_3$): δ 170.0, 166.7, 166.3 (3 × C=O), 133.3, 133.2, 129.8, 129.7, 129.6, 128.5 (Ph), 99.2 (C-1), 69.6 (C-3), 66.2 (C-6), 65.8 (C-5), 55.2 (OMe), 52.1 (C-2), 33.1 (C-4), 23.4 (NHCO*C*$H_3$). LRMS: *m/z* = 427.96 $[M+H]^+$, 450.16 $[M+Na]^+$.

**Methyl 2-acetamido-2,4-dideoxy-α-D-*xylo*-hexopyranoside (6)**

Dibenzoate **5** (1.91 g, 4.47 mmol) was dissolved in anhydrous MeOH (30 mL) and KOH (4.26 g, 76 mmol) was added. The resulting solution was stirred at r.t. for 20 h. The reaction mixture was then neutralized with Amberlite® IR120 ($H^+$), pre-washed with MeOH (3 × 20 mL) in small portions to pH 7. The mixture was filtered, and concentrated to dryness. The residue was then purified by flash chromatography ($CHCl_3$/ MeOH, 4:1) to give the diol **6** as a white solid (886 mg, 90%); m.p. 148 – 150 °C (lit. [2] 153 – 155 °C [MeOH/EtOAc]); $[\alpha]_D$ +157.8 (*c* 0.695, MeOH; lit [2] +186.7); $R_f$ = 0.47 ($CHCl_3$/ MeOH, 4:1); $^1$H NMR (500 MHz, $CD_3OD$): δ 4.62 (d, 1H, $J_{1,2}$ = 3.4 Hz, H-1), 3.81 – 3.71 (m, 3H, H-2, H-3, H-5), 3.48 (app d, 2H, $J$ = 5 Hz, H-6a, 6b), 3.28 (s, 3H, OMe), 1.91 (s, 3H, Ac), 1.92-1.88 (m, 1H, H-4eq), 1.34 (ddd, 1H, $J_{3,4ax}$ = $J_{4ax,5}$ = $J_{4ax,4eq}$ = 12.3 Hz, H-4ax); $^{13}$C NMR (125 MHz, $CD_3OD$): δ 173.8 (C=O), 100.4 (C-1), 69.9 (C-5), 67.1 (C-3), 65.7 (C-6), 56.9 (C-2), 55.4 (OMe), 37.2 (C-4), 22.6 (NHCO*C*$H_3$). LRMS: *m/z* = 219.84 $[M+H]^+$, 242.08 $[M+Na]^+$.

**Methyl 2-acetamido-3,6-di-*O*-acetyl-2,4-dideoxy-α-D-*xylo*-hexopyranoside (7)**

Diol **6** (14 mg, 60 μmol) was dissolved in anhydrous pyridine (2 mL). Acetic anhydride (123 μL, 1.3 mmol) and a catalytic amount of DMAP were added. The reaction mixture was stirred at r.t. for 24 h. MeOH (2 mL) was added at 0 °C to quench access acetic anhydride and the solution was then allowed to stir at 0 °C for 1 h. The mixture was concentrated, and the residue diluted with $CHCl_3$ and washed with 1M HCl, sat. aq. $NaHCO_3$ and brine, dried ($MgSO_4$), filtered and concentrated to give the diacetate **7** as a solid (19 mg, 100%), $R_f$ = 0.25 (4:1 EtOAc/*n*-hexane), m.p. 158 – 160 °C (lit.[2] 158 – 160 °C [hexanes/EtOAc]); $[\alpha]_D$ +77.6 (*c* 0.235, $CHCl_3$; lit.[2] +88.8). $^1$H NMR (500 MHz, $CDCl_3$): δ 5.63 (d, 1H, $J_{2,NH}$ = 9.7 Hz, NH), 5.11 (ddd, 1H, $J_{2,3}$ = 10.0 Hz, $J_{3,4a}$ = 5.1 Hz, $J_{3,4b}$ = 12.0 Hz, H-3), 4.72 (d, 1H, $J_{1,2}$ = 3.5 Hz, H-1), 4.16 (ddd, 1H, H-2), 4.11 (app d, 2H, $J$ = 4.8 Hz, H-6a, 6b), 4.02 – 3.96 (m, 1H, H-5), 3.36 (s, 3H, OMe), 1.98 (ddd, 1H, $J_{3,4eq}$ = 5.1 Hz, $J_{4eq,5}$ = 2.2 Hz, $J_{4eq,4ax}$ = 12.3 Hz, H-4eq), 2.07 (s, 3H, Ac), 2.01 (s, 3H, Ac), 1.94 (s, 3H, Ac), 1.65 (ddd, 1H, $J_{3,4ax}$ = 12.0 Hz, $J_{4ax,5}$ = 12.0 Hz, $J_{4ax,4eq}$ = 12.3 Hz, H-4b); $^{13}$C NMR (125 MHz, $CDCl_3$): δ 171.1, 170.7, 170.0 (3 × C=O), 99.0 (C-1), 68.4 (C-3), 65.6 (C-6), 65.5 (C-5), 55.0 (OMe), 52.2 (C-2), 32.6 (C-4), 23.2, 20.9, 20.7 (3 × Ac). LRMS: *m/z* = 303.88 $[M+H]^+$, 326.04 $[M+Na]^+$.

**2-acetamido-1,3,6-tri-*O*-acetyl-2,4-dideoxy-D-*xylo*-hexopyranose (8)**

A small drop of conc. $H_2SO_4$ was added to a mixture of **7** (42 mg, 140 μmol) and acetic anhydride (50 μL, 3.7 eq.). The mixture was stirred at r.t. for 4 h and then neutralised with sodium acetate to pH 7. The mixture was filtered through a plug of celite and washed with DCM (10 × 15 mL). The combined filtrate and washings were evaporated under reduced pressure to give a pale yellow syrup. The crude product was purified by flash chromatography (EtOAc/toluene, 8:1) to give the triacetate **8** as a white solid (42 mg, 91%). The ratio of α:β anomers was 1.4:1 by $^1$H NMR spectroscopy. α:β anomers : m.p. 96 – 97 °C.

α-**8**: $R_f$ = 0.34 (EtOAc/*n*-hexane, 9: 1); $^1$H NMR (500 MHz, $CDCl_3$): δ 6.17 (d, 1H, $J_{1,2}$ = 3.5 Hz, H-1), 5.55 (d, 1H, $J_{2,NH}$ = 9.0 Hz, NH), 5.18 (ddd, 1H, $J_{2,3}$ = $J_{3,4ax}$ = 11.3 Hz, $J_{3,4eq}$ = 4.9 Hz, H-3), 4.31 (ddd, 1H, H-2), 4.15 – 4.07 (m, 3H, H-5, H-6a, H-6b), 2.14 (s, 3H, Ac), 2.08 – 2.02 (m, 1H, H-4eq), 2.06 (s, 3H, Ac), 2.05 (s, 3H, Ac), 1.93 (s, 3H, Ac), 1.80 – 1.65 (m, 1H, H-4ax); $^{13}$C NMR (125 MHz, $CDCl_3$): δ 171.5, 170.6, 170.1, 168.9 (C=O × 4), 91.8 (C-1), 67.7 (C-5), 67.6 (C-3), 65.2 (C-6), 51.6 (C-2), 32.5 (C-4), 23.1, 21.0, 20.9, 20.7 (Ac × 4); β-**8**: $R_f$ = 0.30 (EtOAc/*n*-hexane, 9: 1); $^1$H NMR (500 MHz, $CDCl_3$): δ 5.59 (d, 1H, $J_{1,2}$ = 8.7 Hz, H-1), 5.57 (d, 1H, $J_{NH}$ = 9.1 Hz, NH), 4.99 (ddd, 1H, $J_{2,3}$ = $J_{3,4ax}$ = 11.3 Hz, $J_{3,4eq}$ = 5.1 Hz, H-3), 4.15 – 4.07 (m, 3H, H-2, H-6a, H6b), 3.87 – 3.82 (m, 1H, H-5), 2.09 (s, 3H, Ac), 2.07 (s, 3H, Ac), 2.04 (s, 3H, Ac), 2.08 – 2.02 (m, 1H, H-4eq), 1.92 (s, 3H, Ac), 1.80 – 1.65 (m, 1H, H-4ax); $^{13}$C NMR (125 MHz, $CDCl_3$): δ 171.0, 170.2, 170.1, 169.7 (C=O × 4), 93.2 (C-1), 70.6 (C-5), 70.1 (C-3), 65.1 (C-6), 53.7 (C-2), 32.5 (C-4), 23.2, 20.9, 20.8, 20.7 (Ac × 4). LRMS: *m/z* = 354.17 $[M+Na]^+$; HRMS: *m/z* calcd for $C_{14}H_{21}NNaO_8$ [M+Na]+: 354.1159; found: 354.1162.
